# Bounded optimality of time investments in rats, mice, and humans

**DOI:** 10.1101/2024.12.09.627552

**Authors:** Torben Ott, Marion Bosc, Joshua I. Sanders, Paul Masset, Adam Kepecs

## Abstract

Time is our scarcest resource. Allocating time optimally presents a universal challenge for all organisms because the future benefits of time investments are uncertain. We developed a normative framework for assessing bounded optimality in time allocation, emphasizing the accuracy of future predictions, independent of subjective costs and benefits. In a common decision task across humans, rats, and mice, we varied uncertainty by titrating ambiguous sensory evidence and measured the time each subject was willing to invest post-decision. We observed that all species and subjects invested more time when they were more likely to be correct, which reflected a statistical confidence of uncertain evidence. Time allocation strategy approached the lower bound of optimality, indicating an accurate decision-by-decision assessment of confidence in the likelihood that waiting will pay off – independent of the subjective payoff values and time costs. We demonstrate that an elementary algorithm based on a drift-diffusion process algorithm can implement this optimal time investment strategy. These results illuminate the computational mechanisms governing rational time investment, showing that humans, rats, and mice can maximize payoffs via confidence-guided time allocation.

**Highlights:**

- Computational and behavioral framework to assess bounded optimality of investments.
- Humans, rats, and mice invest more time to obtain more likely payoffs, in proportion to statistical confidence.
- Time investment was close to optimal model predictions, reflecting bounded optimality of investments under uncertainty.
- Bounded-optimal time investment may be an evolutionary ancient adaptive behavioral strategy.

## Introduction

Time is a universal currency that all organisms, from the tiniest microbe to the largest mammal, must use wisely to thrive. Humans and other animals alike engage in the deliberate allocation of time to various pursuits, including learning new skills, foraging for food, and leisurely activities. Classic economic theories propose how rational investments should balance the investment’s expected gain against the cost of forgoing other opportunities (Becker 1965; Keynes 1936). Yet, when faced with uncertainty, a plethora of studies have demonstrated that human behavior often veers from these rational decision models (Kahneman and Tversky 1972; 1979; Camerer 1999), constrained by limited cognitive capacities and environmental resources (Simon 1997; Gigerenzer and Todd 1999; Sent 2018). Despite time’s status as a universal currency, it is often unclear if humans and other animals make optimal use of their time. Here, we investigated time investment strategies in different species to shed light on the evolutionary origin of rational, and irrational, investment decisions.

The study of rational time investments has been hindered by two key obstacles. First, we lack a common behavioral framework to assess time investments. Most human studies rely on hypothetical choices lacking experiential components, whereas real-life investment decisions require learning from past experiences and may not show the same cognitive biases, a discrepancy known as the description-experience gap (Basile, Fabien, and Stefano 2020; Hertwig and Erev 2009). Further, such description-based studies cannot be translated to animals that use tangible outcome: value-based decision-making (Wikenheiser, Stephens, and Redish 2013), delay discounting (Frost and McNaughton 2017; Hayden 2016), and decision confidence (Lak et al. 2014; Masset et al. 2020; Stolyarova et al. 2019).

A second challenge to study time investments is the lack of a theoretical framework given that the costs and benefits can be highly subjective and idiosyncratic. There is no universal formula for optimal time allocation without assumption about the value, or utility, of time and rewards (Sent 2018; Houston and Rosenström 2023). For instance, browsing the internet might be wasting time for one individual while maximizing the happiness of another. For these reasons, defining an optimal investment strategy often relies on assumptions about the shape of the agent’s utility functions, which is typically unknown. Rational investment decisions also require predicting the investment’s likely outcome (Becker 1965; Sen 2020). When confronted with uncertainty, humans and animals often struggle to accurately assess risks and payoffs, distorting their probability estimates (Camerer 1999; Tversky and Kahneman 1981; Constantinople, Piet, and Brody 2019; Houston and Rosenström 2023; Yang, Li, and Stuphorn 2022; Stauffer et al. 2015). The absence of a computational and behavioral framework that addresses both challenges impedes normative interpretations of time investment behavior.

To address these issues, we developed a framework to comparatively assess the ‘bounded optimality’ of time investments across species, defined as deviations from optimal time allocation of a utility-maximizing agent. Behaviorally, we used a post-decision wagering task in which subjects first make a binary choice about ambiguous auditory stimuli before placing a bet on their decision by investing time continuously until giving up and moving to the next decision. Importantly, the same task contingencies could be used across species: in humans as a video game and in mice and rats working for water rewards. We developed a normative investment model based on Keynesian investment theory and found that humans, rats, and mice invested time in proportion to their degree of decision confidence. All species invested time close to optimal model predictions under arbitrary subjective utility functions, thus demonstrating bounded optimality with respect to decision confidence to optimize investment decisions. We found that humans and other animals are smart intuitive investors, who leverage their sense of confidence to maximize future gains.

## Results

### Variable time investment task

To quantitively assess time investment behavior in humans, rats, and mice we adopted a previous task design (Lak et al. 2014; Masset et al. 2020) with three key features: (i) self-controlled variable time investment, (ii) experimenter-controlled uncertainty in evidence, (iii) cross-species experiential task design. Subjects invested variable time into decisions based on ambiguous auditory evidence, which allowed us to systematically vary the amount of evidence for a decision and simultaneously observe each subject’s choice and time investment behavior. Task design, auditory stimuli, and time investment were analogous across species (Figure 1A-B): Humans, rats, or mice self-initiated a trial, which triggered delivery of an auditory stimulus. Auditory stimuli were identical for all species and consisted of binaural streams of Poisson-distributed clicks with different underlying click rates between the left and right click stream. In a series of consecutive decisions (‘trials’), subjects had to determine the side with the higher underlying click rate and give the correct response (left or right) to receive a money reward (humans, Figure 1A) or water reward (rats and mice, Figure 1B). The strength of evidence for each decision was given by the difference in the number of clicks between left and right divided by the total number of clicks (‘binaural contrast’). Human subjects performed a gamified version of this task, for which pilot experiments showed improved choice and investment behavior (Figure 1C; cf. Methods).

**Figure 1.**
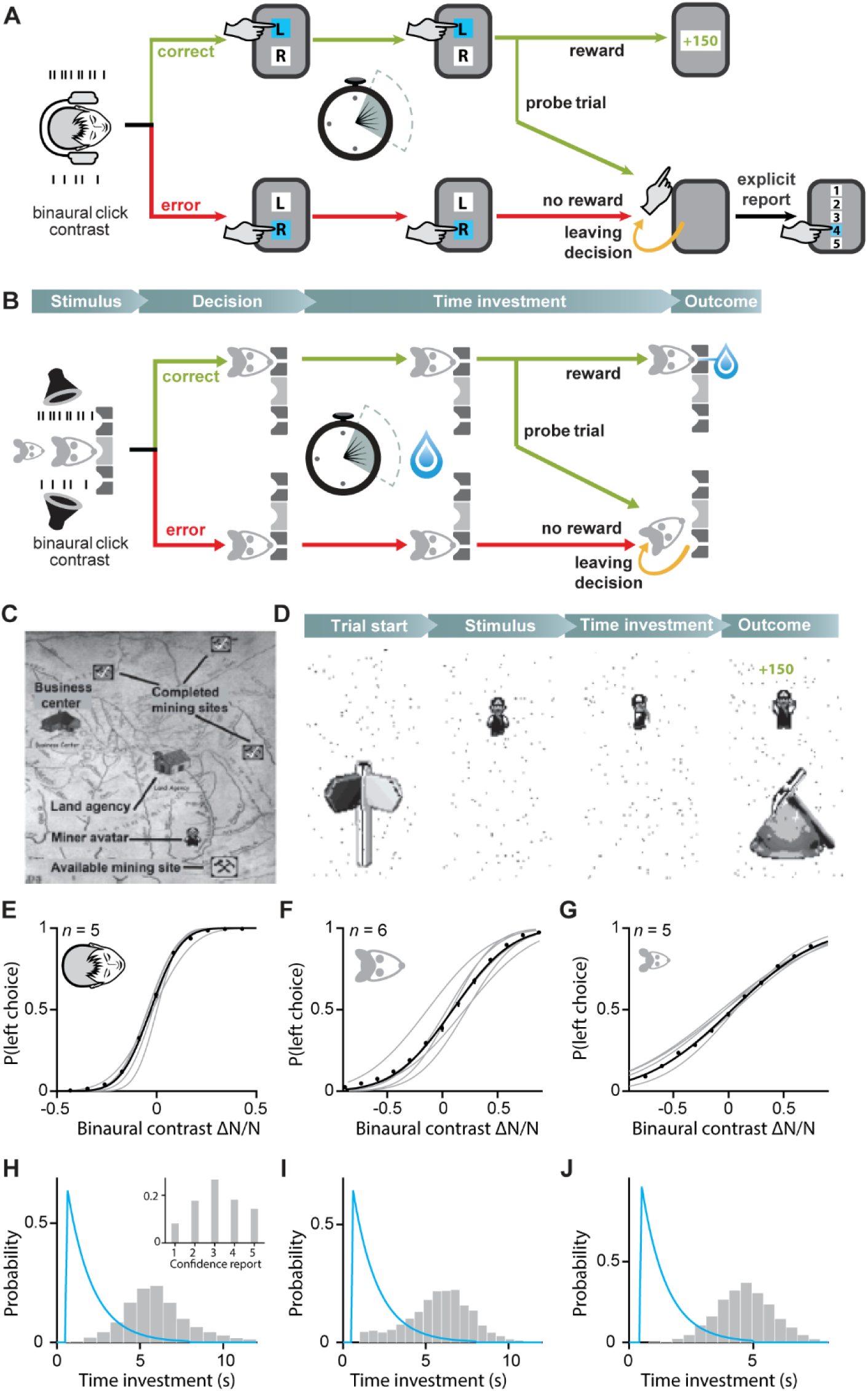
Post-decision time investment task for humans, rats, and mice. **(A)** Time investment task in humans. Humans discriminated binaural stream of clicks and indicated their choice with a button press (left or right) in a series of consecutive trials. Reward delivery (credits) was randomly delayed, and no feedback was given on error trials and a random subset of correct trials (probe trials, 10 %). We thus obtained choice and time investment in error and probe trials. In probe trials and a subset of error trials (30 %) we asked subjects to explicitly report their confidence on a 5-point scale. **(B)** Analogous time investment task in rats and mice. Rats or mice discriminated between the same binaural click streams as humans and indicated their choice by entering a choice port (left or right), and reward delivery (water) was randomly delayed. As in humans, we obtained choice and time investment in probe (10 % trials in rats, 5 % in mice) and error trials, in which no reward or other feedback was given. **(C)** In humans, we gamified this task. In brief, humans controlled a miner avatar and could enter mining sites on a map to initialize a 5-minute block of trials. Following a trial block, earned credits for correct choices could be exchanged for candy in the business center. **(D)** After entering a mining site, trial start was indicated by appearance of a sign post, followed by the auditory stimulus. The choice (left or right) was indicated by the avatar facing left or right and the background moving in opposite direction for as long as the subject held the choice button. Visual feedback indicated reward delivery and credits added to the account. **(E)** Psychometric curve for human subjects showing average choices to the left as a function of evidence strength, defined as the difference in the number of clicks between left and right divided by their sum. Gray lines show single subjects, black line shows pooled subject data. Error bars indicate 95 %-confidence intervals. **(F)** Psychometric curves for rat subjects, conventions as in E. **(G)** Psychometric curves for mouse subjects, conventions as in E. **(H)** Time investment distribution in probe and error trials (pooled across subjects) (gray) and reward delay distribution (blue line). Inset shows explicit confidence reward distribution. **(I)** Time investment distribution for rats (pooled across subjects). **(J)** Time investment distribution for mice (pooled across subjects).

All subjects’ decisions were explained by the strength of auditory evidence with perceptual noise. For easy decisions with high binaural contrast, subjects made few mistakes (i.e., showed low lapse rates) and subjects showed graded accuracy in more difficult decisions (Figure 1E-G). Subjects’ choices were well predicted by a generative model with Gaussian noise, which we fit with cumulative Gaussian distribution to each subject’s choice behavior (Figure 1E-G, Table 1, p < 0.001 for each subject against shuffled data, permutation test). Together, the combination of evidence strength and choice quantitatively determines the subjects’ choice uncertainty. Subjects were incentivized to invest variable amounts of time in their choices. After committing to a choice, reward delivery for correct choices was randomly delayed (humans and rats: 0.5–8 s, exponentially distributed with decay constant *τ* = 1.5; mice: 0.5–5 s, *τ* = 1.0 s). In error trials and a subset of correct trials (probe trials, humans and rats: 10 %; mice: 5 %), no reward or feedback was given. Thus, subjects waited for a self-determined amount of time, before giving up waiting and moving to the next decision. The subjects’ waiting time in a trial constitutes a graded time investment into their decision. Since access to water was restricted to behavioral sessions of 2-3 hours per day in rats and mice, rodent subjects were incentivized to adjust their time investment behavior to maximize water reward intake. In humans, incentive structures in psychophysical tasks are more complex. We therefore sought to (i) create strong incentive structures, and (ii), promote task engagement by gamifying the task in humans. In this gamified task, using identical sensory stimuli and trial contingencies, subjects used virtual ‘Geiger counters’ to locate (left or right) ‘plutonium sites’ in ‘mines’. ‘Mining expeditions’ were limited to brief 5-minute blocks incentivizing adaptive time allocation, and 15 blocks per session. Reward credits were awarded for each correct choice (150 credits per correct choice), and could be redeemed for different types of candy after the session was complete. High performance was also incentivized with bonus payments. Thus, optimizing time investment earned the subjects more money. Overall, we observed variable time investment behavior across trials (Figure 1H-J; humans: 5.9 [4.8–7.0] s (median [IQR]); rats: 6.3 [4.9–7.2] s; mice: 4.8 [3.9–5.2] s, data pooled across subjects).

**Table 1.**
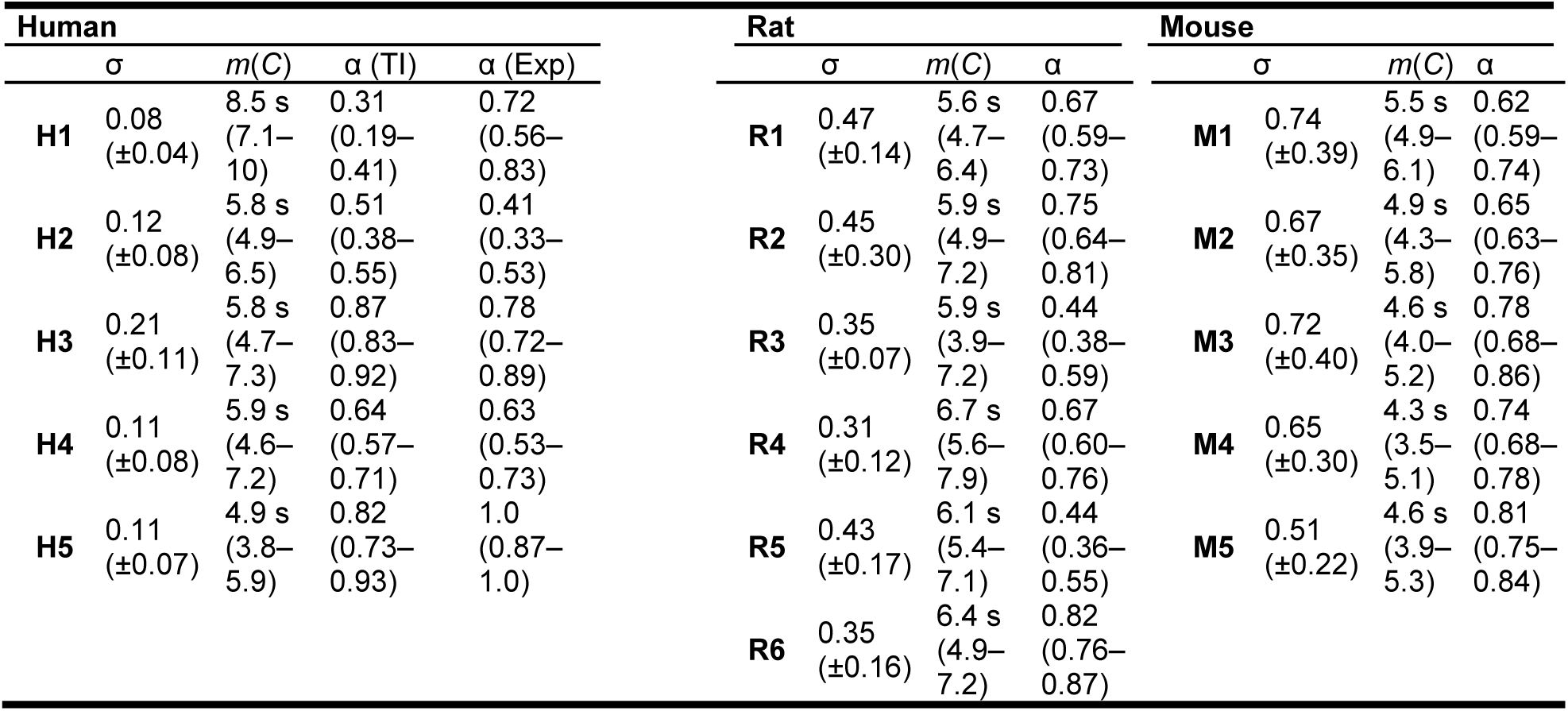
Confidence-guided time investment model results for each subject. For perceptual noise σ, values indicate standard deviation of a cumulative Gaussian fit to psychometric function (95 %-confidence intervals, *t*-statistic). To describe the mapping function *m*(*C*), values indicate median time investment (inter-quartile range). For investment efficiency α, values indicate maximum-likelihood fit (95 %-confidence interval, bootstrap). TI, time investment; Exp, explicit reports.

### Bounded optimality of confidence-guided time investment

What constitutes an optimal investment strategy in this task, that is, how should one invest time to maximize overall reward? Our aim is to assess the theoretical ideal strategy for time allocation based on uncertain evidence that maximizes subjective payoff, which we call ‘bounded optimality’. This approach focuses on objective effectiveness of time allocation, while allowing for arbitrary subjective factors such as the perceived value, or utility, of payoffs and cost of time, which are inherently individual and challenging to measure. Drawing on the principles of rational task analysis (Neth, Sims, and Gray 2016), we derive the optimal time investment strategy for our task. This analysis will enable us to delineate the efficiency of time investment behavior from ‘random and ‘optimal’ performance. i.e., the bounded optimality of time investment.

According to Keynesian investment theory, an investment is deemed favorable if the expected future return exceeds its opportunity cost (Becker 1965; Keynes 1936). In this context, the expected return *R* is the amount of water or monetary reward, weighed by the probability of receiving a reward after investing time *T*. The opportunity cost *K* of a time investment is the amount of reward, on average, that we forgo by waiting for a reward instead of moving on to other investment opportunities, i.e., moving on to the next trial in our task. The value of time investment *T, V(T),* is then the difference between its expected return *R(T)* and opportunity cost *K(T)*, reflecting the potential missed gains by foregoing other investments:

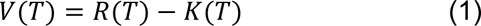

Optimal time investment *T_opt_* maximizes *V(T)*,

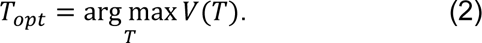

In this variable time investment task, reward magnitude was fixed, and the expected return in a trial is thus determined by the investor’s time-varying reward expectancy: the probability that reward arrives in the next time interval from *t* to d*t* after waiting time *t* without receiving a reward. The expected return *R(T)* for time investment *T* is then given by integrating the reward expectancy per unit time, defined as the hazard rate of reward ρ(*t*):

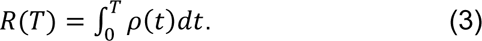

We assumed a stable opportunity cost, represented by κ, for each unit of time invested, based on the premise that the subjects’ behavioral strategies and their subjective valuation of rewards remained unchanged. This assumption is deemed reasonable because the task did not involve any alterations in stimulus and reward conditions, and can be verified by analyzing the data for any significant deviations or non-stationarities in behavior or performance over time. The investment’s opportunity cost then is *K(T)* = *κT* and with (1) and (3) the value of time investment *V(T)* is given by

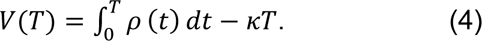

An investor’s challenge is to maximize the value function *V* with respect to time investment *T.* Critically, the hazard rate of reward, *ρ(t),* depends on (i) the investor’s decision confidence *C*, the probability that the current trial’s choice will be rewarded, and on (ii) the experimenter-defined temporal reward distribution during the waiting time period, *P_rew_(t)* (see Methods for details). We chose an exponential reward distribution *P_rew_(t)*, which implies a flat (constant) hazard rate of reward ρ(*t*) when C = 1. For C < 1, the hazard rate of reward ρ(*t*) is monotonically decreasing. Therefore, the value function *V(T)* is concave and maximal at

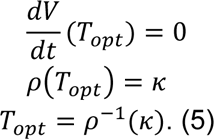

Since the investor’s confidence varied in each trial, and we treat the opportunity cost as stable, we express optimal time investment in each trial as a function of confidence, i.e.

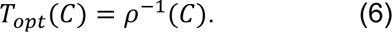

This result shows that optimal time investment should reflect decision confidence, the estimated probability that a decision will lead to a payoff. Here, optimal time investment refers to the statistically optimal use of the subjective percept of the sensory evidence delivered in our task for time allocation. We leveraged the relationship between confidence and time investment to develop a generative optimal time investment model (Figure 2A).

**Figure 2.**
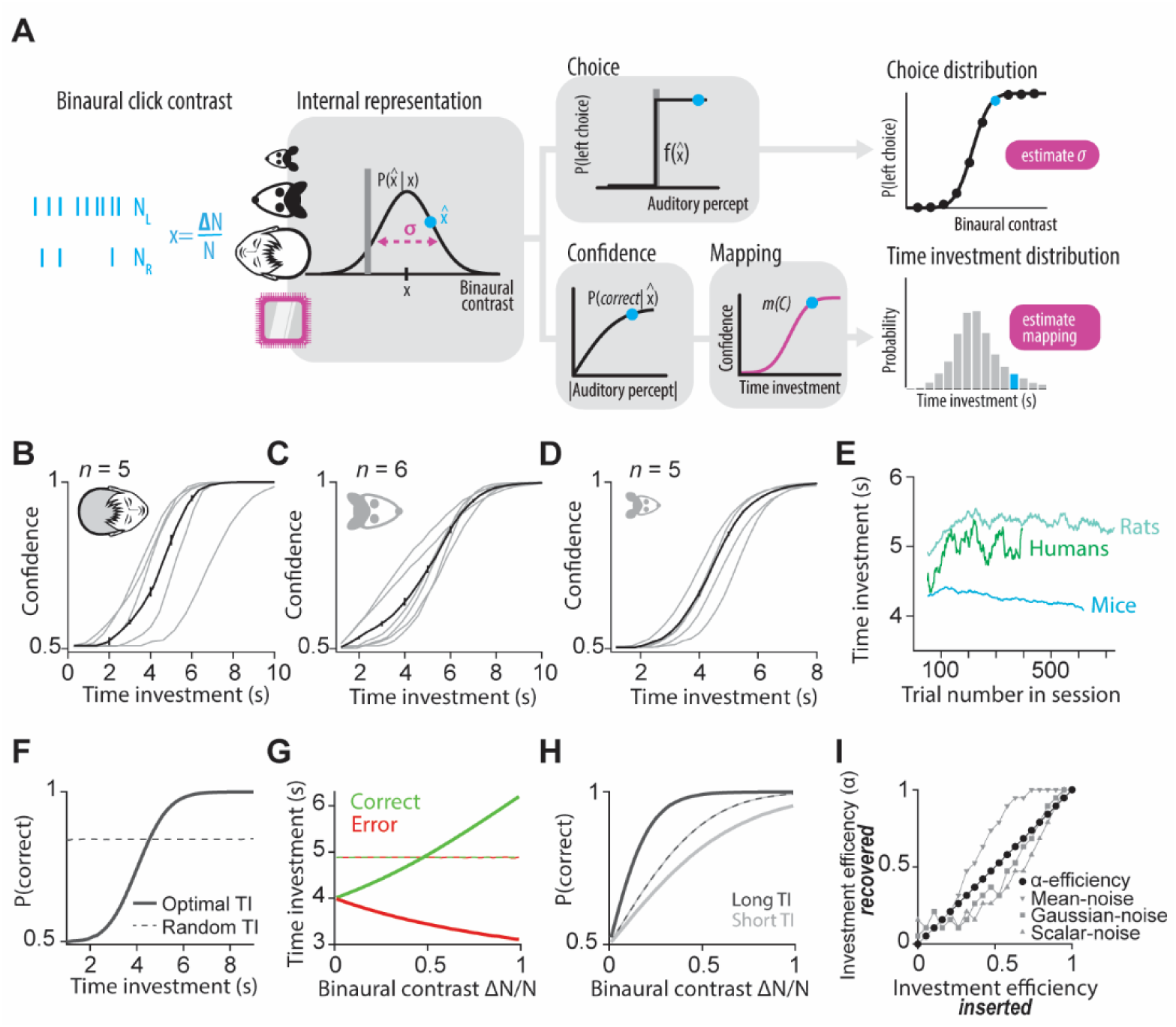
Optimal time investment model to quantify bounded optimality of investment behavior. **(A)** A model for optimal time investment guided by decision confidence. See text and Methods for details. N_L_, left clicks; N_R_, right clicks; *x*, evidence strength (difference in clicks ΔN = N_L_ - N_R_ divided by the sum N = N_L_ + N_R_); *x̂*, percept (internal representation of *x*); σ, perceptual noise; *C*, confidence; *m*(*C*), mapping function. **(B)** Estimated mapping function *T* = *m*(*C*) between confidence *C* and time investment *T* for human subjects. Gray lines, individual subjects; black line, pooled subject data. Error bars indicate 95 %-confidence intervals. **(C)** Estimated mapping functions m(C) for rat subjects, conventions as in B. **(D)** Estimated mapping functions for mouse subjects, conventions as in B. **(E)** Time investment across trials within behavioral sessions was mostly stable (50 trial running average). **(F)**–**(H)** Model-predicted optimal time investment as a function of task performance and evidence (based on σ and *m*(*C*) of a single human subject). **(F)** Model-predicted calibration curve for optimal investment (solid line) and random investment not related to confidence (dashed line, shuffled time investment across trials). **(G)** Model-predicted vevaiometric curve, conventions as in E. **(H)** Model-predicted conditioned psychometric curves, conventions as in E. **(I)** Model recovery test for simulated data across different noise models. α-efficiency captures the noise imputed by different noise models. Values lower than 1 correspond to sub-optimal investment behavior, whereas values larger than 0 indicate that investment behavior is guided by confidence.

A key feature of our decision tasks is that trial-to-trial variations of confidence can be readily inferred from the observed choice behavior. We use a generative decision model to estimate the subjects’ perceptual noise *σ* by fitting a truncated Gaussian distribution to individual choice behavior (Figure 2A, top row, cf. Figure 1E-G, Table 1). The standard deviation *σ* of this distribution provides an estimate of the perceptual noise. Under the assumption of Gaussian perceptual noise, we inferred the distribution of confidence for a given strength of evidence and choice using a normative statistical model of confidence. In other words, we generated percepts *x̂* (here, a percept is an internal representation of sensory evidence *x*) based on evidence *x* with Gaussian perceptual noise σ and calculated the degree of confidence *C* = *P(correct|x̂,choice)* associated with each percept (Figure 2A center). Next, we converted each confidence estimate to a predicted optimal time investment (Figure 2A, bottom row).

To predict the optimal time investment, we need to convert decision confidence (an estimated probability) into seconds of time investment, using a mapping function *T = m(C)*. According to equation (6), the mapping function from confidence to optimal time investment has a theoretical solution. Specifically, the theoretical mapping function *m(C)* is the inverse hazard rate of reward, *m(C)* = ρ^−1^(*C*) (see Methods; Lak et al., 2014). However, the validity of this theoretical solution relies on several strong assumptions. First, the subjective hazard rate of reward is likely to deviate from its theoretical form determined by *P_rew_(t)* (see Methods) due to uncertainty in time estimates. The variance of time estimation increases linearly with time, called scalar timing (Gibbon 1977; Malapani and Fairhurst 2002). Second, we cannot be certain about the precision with which the investor learned the reward distribution *P_rew_(t)*, from which we estimate the hazard rate. Thirdly, we do not know the investor’s utility function with respect to earned reward or effort spent waiting. Finally, we do not know possible interactions between time and utility, that is, the investor’s value of time. Due to these uncertainties, we sought to empirically determine the mapping function between confidence and time investment, which would allow us to isolate the contribution of decision confidence to time investment behavior in a utility-maximizing agent. To infer the mapping function *T = m(C)* between confidence *C* and time investment *T* we assume that *m* is a monotonically increasing function of *C*, and that the mapping function is stable across time (Figure 2E). In other words, longer time investments reflect higher confidence. We can then use the empirically observed time investment distribution *F_T_*, and the model-predicted cumulative confidence distribution, *F_C_*, to define *m(C) = F_T_^-1^(F_C_(C))* (see Methods) (Figure 2B-D). This mapping function from confidence level to time investment duration is fully constrained by the empirical time investment distribution, while being agnostic about how subjects adjusted time investment based on confidence in individual trials. Notably, if we randomly reorder the time investment data across trials—thus disconnecting actual time investments from their corresponding confidence levels—we would still derive the same mapping function *m(C)*. In this way, our model predicts optimal time investment guided by confidence for each trial *k*, with *T*_opt,k_ *= m(C_k_)*, producing a statistically optimal allocation of time based on the available evidence, without additional free parameters and allowing for arbitrary (monotonic) utility functions of confidence against time.

Together, the key ingredients of our optimal time investment model are (i) inferred perceptual noise from the psychometric choice behavior, (ii) inferred statistical decision confidence, and (iii) estimated mapping function between confidence and time investment (Figure 2A). This model is well-suited to make several testable predictions about optimal time investment in the observed time investment patterns across species. First, optimal time investment predicts the average accuracy of a decision-maker, thus yielding a confidence calibration curve (Figure 2F). Second, optimal time investment is positively correlated with evidence strength for correct choices, but negatively correlated with evidence strength for error choices, thus predicting the outcome of a decision even for the same evidence strength (vevaiometric curve, from Greek *vevaios*, certain) (Figure 2G). Finally, the conditioned psychometric curve demonstrates that for high-investment decisions, average accuracy is higher even for the same evidence strength in comparison to low-investment decisions (Figure 2H). These three signatures serve as qualitative benchmarks for confidence-guided time investment (Sanders, Hangya, and Kepecs 2016; Hangya, Sanders, and Kepecs 2016).

To quantify deviations from optimal time investment, i.e., bounded optimality of time investment with respect to leveraging confidence, we introduced a single free parameter (α: investment efficiency) that summarizes diverse sources of noise. An α value of 1 means perfect efficiency, while 0 produces randomly resampled time investments across all decisions (i.e., corresponding to shuffling all time investment data across trials). This investment efficiency parameter thus captures potentially diverse sources of noise such as ‘metacognitive noise’, i.e., deteriorated confidence estimates, timing uncertainty, or other sources of variability in investment behavior not driven by confidence. We used Monte-Carlo models and maximum-likelihood fitting to estimate investment efficiency. Model-fitting recovered noise-infused simulated data, generated by simulating trials with noise from a range of different noise models (Figure 2I). Note that investment efficiency, *α,* provides a lower bound to optimal use of decision confidence. The key reason is that investment efficiency, *α,* captures not only sources of variability with respect to decision confidence (‘metacognitive noise’), but also variability of the mapping function *m(C)*. Moment to moment or session to session variations of temporal estimation, effort cost, or utility functions will be captured by variability in the mapping function. Thus, assuming a stable mapping function across trials and sessions leads to an underestimate, a lower bound, on the investment efficiency.

### Humans, rats, and mice optimize investment decisions guided by decision confidence

The confidence-guided time investment model allowed us to predict optimal time investments without free parameters and calculate investment efficiency for each subject in humans, rats, and mice. Across all species, the observed time investment behavior followed optimal model predictions. To quantify this we examined the expected time investment patterns for key behavioral signatures of confidence (dashed lines, Figure 3A-C, Figure 4A-C, Figure 5A-C): First, time investment predicted accuracy across the entire range of accuracies (about 50 %–100 %), demonstrating that time investments cover the full range of possible confidence levels and is thus well-calibrated (human single subject example (Figure 3A), *ρ* = 0.87 (Spearman rank correlation), p < 0.001; human pooled subjects (Figure 3E), *ρ* = 0.85, p < 0.001; rat single subject example (Figure 4A), *ρ* = 0.71, p = 0.02; rat combined subjects (Figure 4E), *ρ* = 0.91, p < 0.001; mouse single subject example (Figure 5A), *ρ* = 0.94 p < 0.001; mouse combined subjects (Figure 5E), *ρ* = 0.92, p < 0.001). Second, time investments were positively related to the strength of evidence (absolute binaural contrast) in correct trials (slope > 0, linear regression, p < 0.001, for each single subject example and pooled subject data) and negatively related in error trials (slope < 0, p < 0.001, linear regression, for each single subject example and pooled subject data) as revealed by the vevaiometric curve for human (Figure 3B,F), rat (Figure 4B,F), and mouse (Figure 5B,F) subjects. Finally, when splitting trials into high and low time investments (split by median time investment, cf. Table 1), the conditioned psychometric function was steeper for high time investment compared to low time investment trials, i.e., time investment predicts accuracy even for the same strength of evidence in human (Figure 3C,G), rat (Figure 4C,G) and mouse (Figure 5C,G) subjects (Δslope > 0, logistic regression, p < 0.01 (bootstrap) for each single subject example and pooled subject data). Thus, subjects from all species adjusted time investments according to their degree of confidence. In many cases, the overall predictions for optimal time investment fell within the observed variation of the investment behavior.

**Figure 3.**
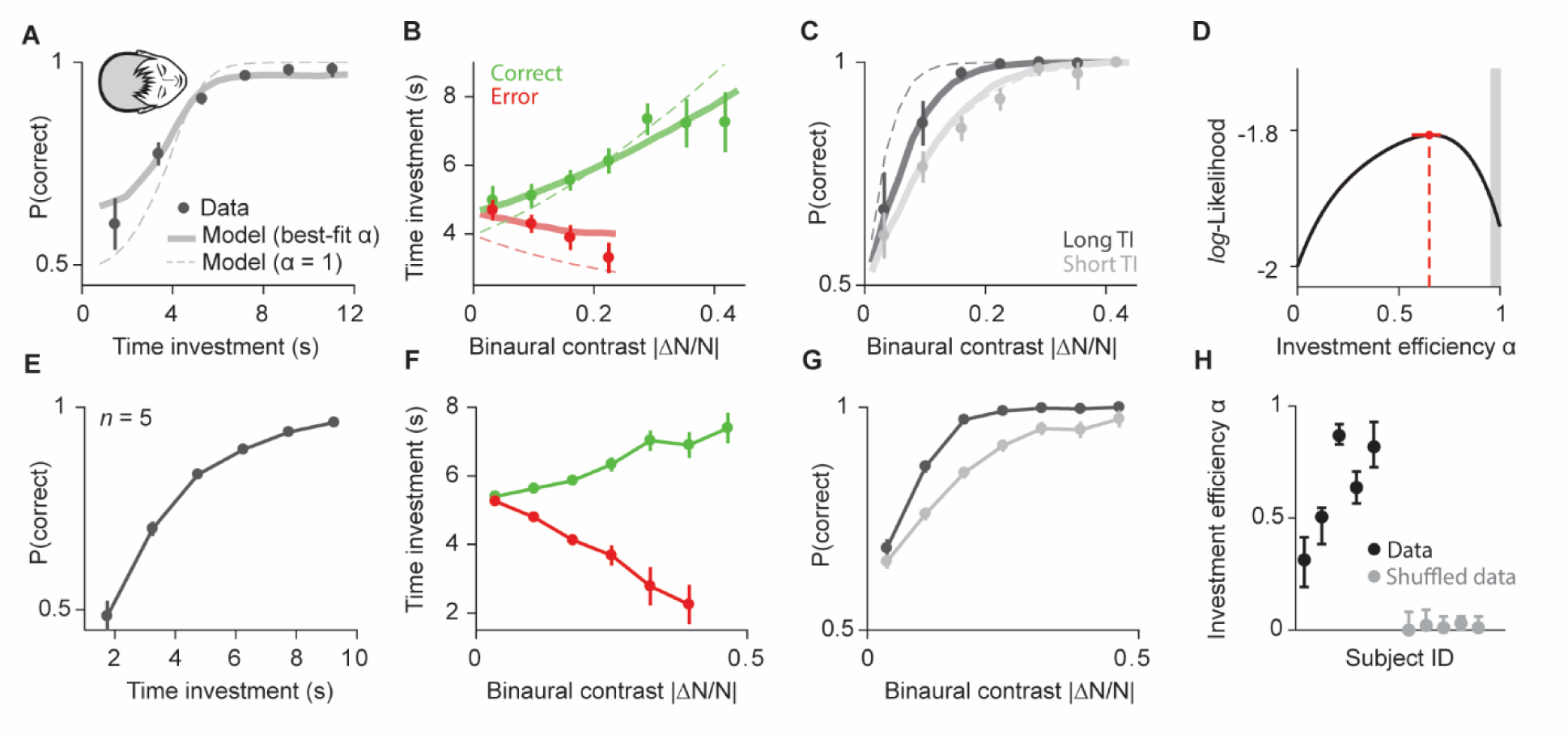
Humans optimized time investment guided by using decision confidence. **(A)–(C)** Model-predicted time investment for optimal (dashed line) and best-fit investment efficiency α (solid line) and observed time investment (points) for single human subject (H4). Observed time investment (TI) showed signatures of statistical decision confidence for calibration curve (A), vevaiometric curve (B), and conditioned psychometric curves (C). **(D)** Likelihood function for same subject maximal at α_best_ = 0.64 (cf. Table 1). **(E)–(G)** Signatures of statistical confidence in pooled human subject data showing calibration curve (E), vevaiometric curve (F), and conditioned psychometric curve (G). **(H)** Best-fit investment efficiency α per human subject (black dots) and shuffled control (gray dots). Error bars indicate 95 %-confidence intervals.

**Figure 4.**
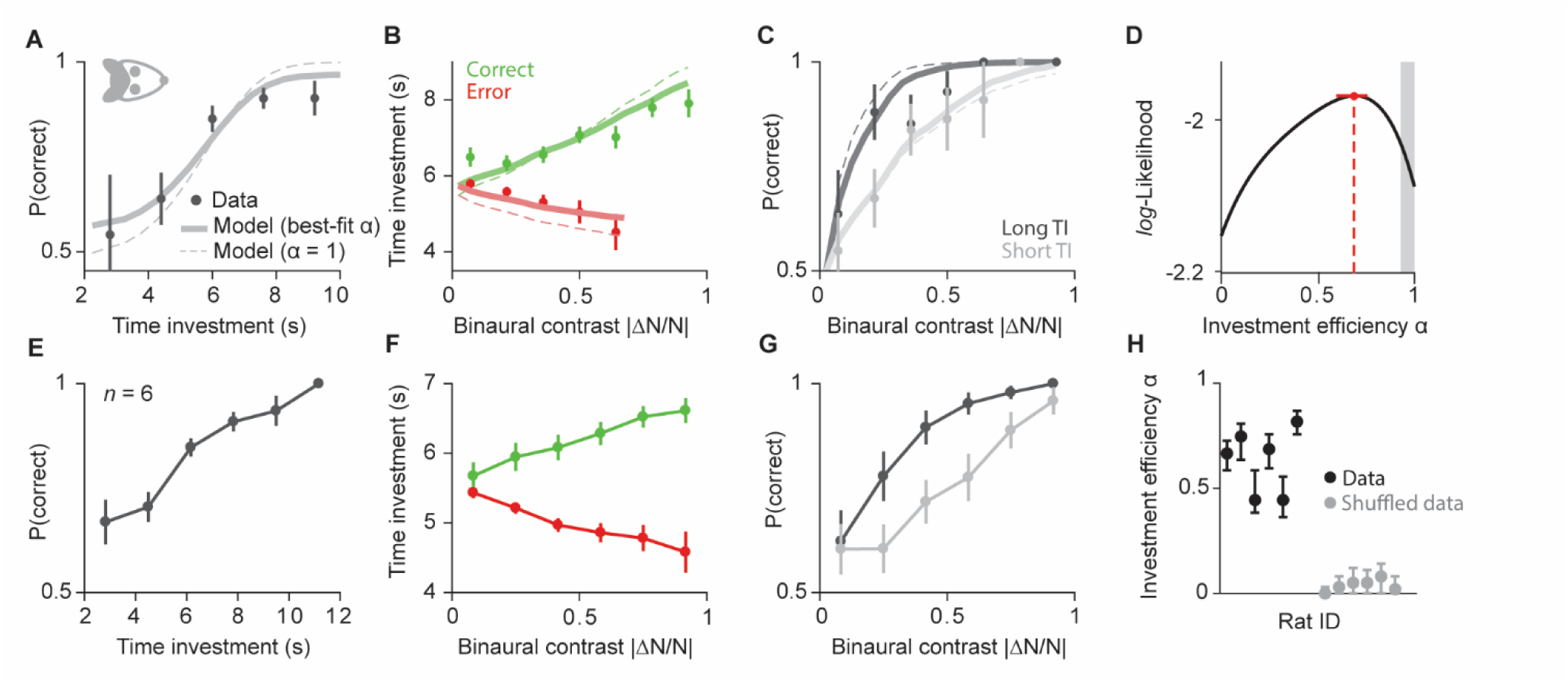
Rats optimized time investment guided by using decision confidence. **(A)–(C)** Model-predicted time investment for optimal (dashed line) and best-fit investment efficiency α (solid line) and observed time investment (points) for single rat subject (R4). Observed time investment (TI) showed signatures of statistical decision confidence for calibration curve (A), vevaiometric curve (B), and conditioned psychometric curves (C). **(D)** Likelihood function for same subject maximal at α_best_ = 0.67 (cf. Table 1). **(E)–(G)** Signatures of statistical confidence in pooled rat subject data showing calibration curve (E), vevaiometric curve (F), and conditioned psychometric curve (G). **(H)** Best-fit investment efficiency α per rat subject (black dots) and shuffled control (gray dots). Error bars indicate 95 %-confidence intervals.

**Figure 5.**
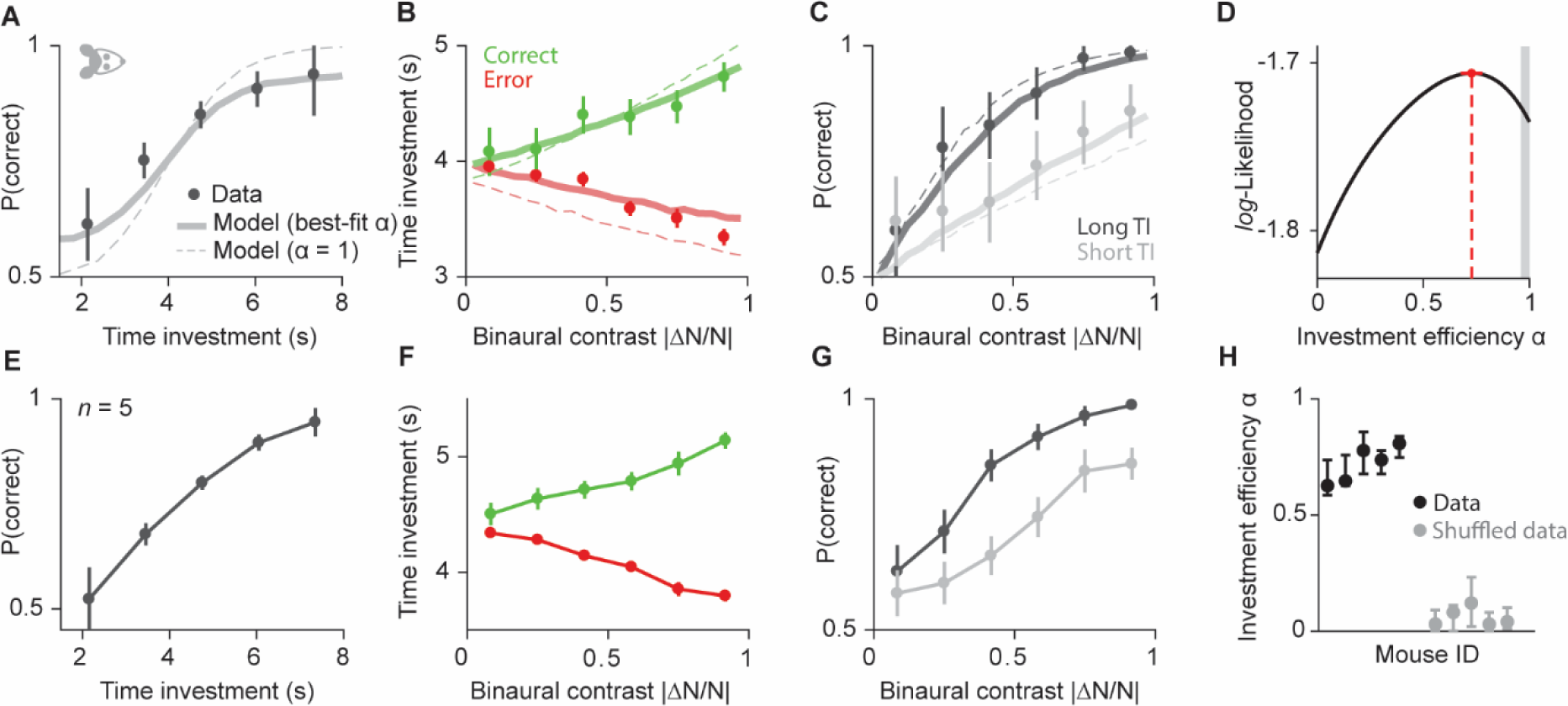
Mice optimized time investment guided by using decision confidence. **(A)–(C)** Model-predicted time investment for optimal (dashed line) and best-fit investment efficiency α (solid line) and observed time investment (points) for single mouse subject (M4). Observed time investment (TI) showed signatures of statistical decision confidence for calibration curve (A), vevaiometric curve (B), and conditioned psychometric curves (C). **(D)** Likelihood function for same subject maximal at α_best_ = 0.74 (cf. Table 1). **(E)–(G)** Signatures of statistical confidence in pooled mouse subject data showing calibration curve (E), vevaiometric curve (F), and conditioned psychometric curve (G). **(H)** Best-fit investment efficiency α per mouse subject (black dots) and shuffled control (gray dots). Error bars indicate 95 %-confidence intervals.

We next measured how close subjects invested time to optimal model predictions by fitting a single free parameter, investment efficiency *α,* to each subject’s time investment behavior. We estimated the model’s likelihood function using Monte-Carlo methods and determined the investment efficiency α that maximized each subject’s likelihood function for human (Figure 3D), rat (Figure 4D), and mouse (Figure 5D) subjects. As expected, time investment predictions from best-fit investment efficiency models yielded improved fits to the data (thick lines, Figure 3A-C, Figure 4A-C, Figure 5A-C). Investment efficiency was larger than zero (corresponding to shuffled time investment unrelated to confidence) for all subjects across all species (α > 0, p < 0.001, permutation test) and, in most cases, investment efficiency was high with α > 0.5 (13/16 subjects), indicating that subjects across all species were efficient investors with low metacognitive noise (Figure 4H, Figure 5H, Figure 6H, cf. Table 1).

**Figure 6.**
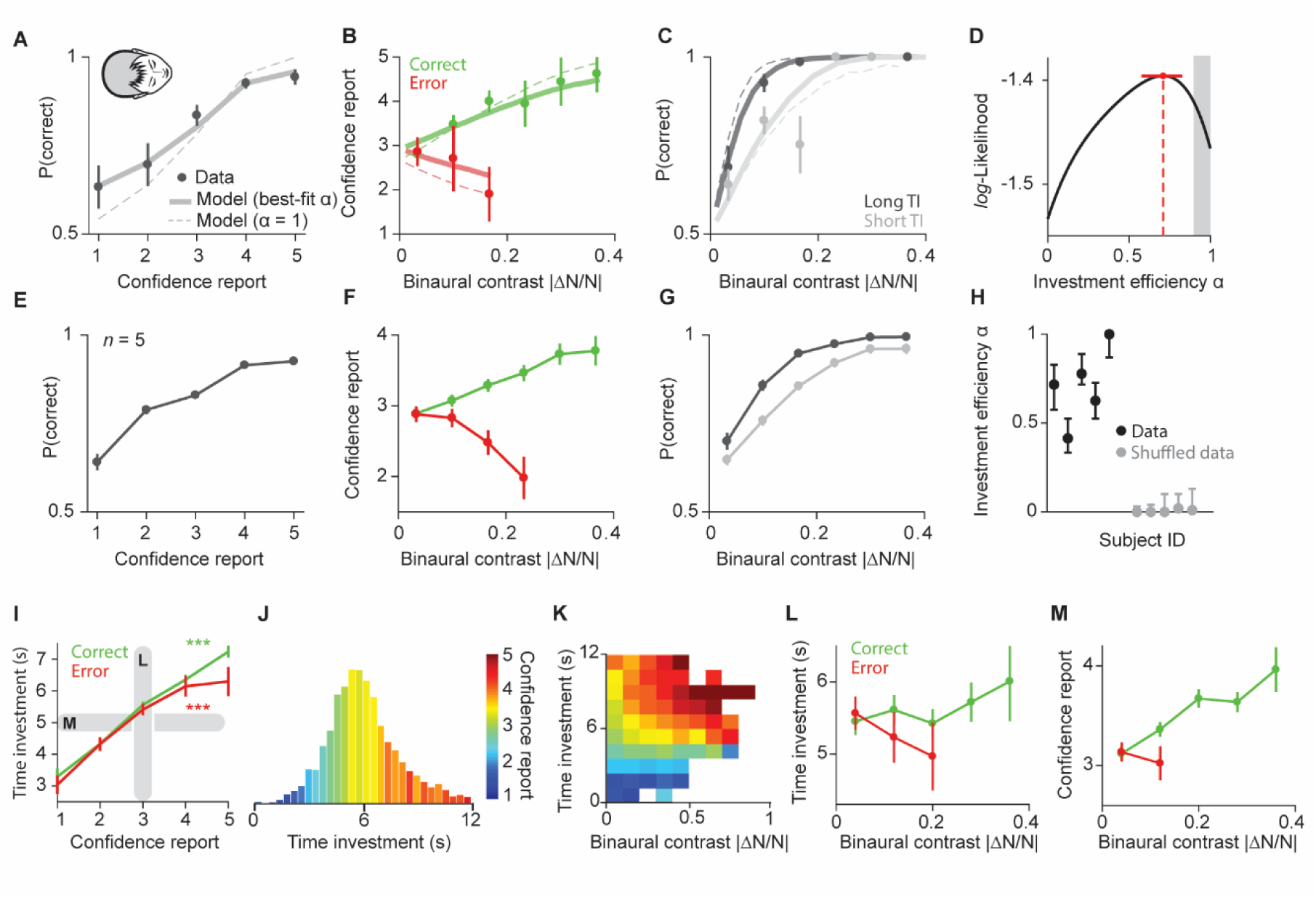
Human explicit confidence reports showed signatures of statistical decision confidence and were predicted by time investment. **(A)–(C)** Model-predicted confidence reports for optimal (dashed line) and best-fit investment efficiency α (solid line) and observed confidence reports (points) for single human subject (H1). Observed confidence reports showed signatures of statistical decision confidence for calibration curve (A), vevaiometric curve (B), and conditioned psychometric curves (C). TI, time investment. **(D)** Likelihood function for same subject maximal at α_best_ = 0.72 (cf. Table 1). **(E)–(G)** Signatures of statistical confidence in pooled human subject data showing calibration curve (E), vevaiometric curve (F), and conditioned psychometric curve (G). **(H)** Best-fit investment efficiency α per human subject (black dots) and shuffled control (gray dots). **(I)** Correlation between time investment and explicit confidence reports for correct and error trials. **(J)** Time investment distribution colored by explicit confidence reports showed strong correlation between the two reports. **(K)** 2D map between time investment and binaural contrast showed that time investment and explicit reports were correlated even for the same evidence. **(L)** Vevaiometric curve for a fixed explicit confidence report of 3. **(M)** Vevaiometric curve for a narrow range of time investment values around the median investment (quantile 0.4–0.6). Error bars indicate 95 %-confidence intervals.

### Human time investment and verbal confidence reports reflect shared and independent confidence information

Humans use both experiential and verbal assessments of their decisions. We could therefore assess how time investment, an experiential or ‘implicit’ report of confidence, relates to declarative or ‘explicit’ verbal reports in human subjects. While previous work demonstrated that verbal confidence reports convey statistical decision confidence estimates (Sanders, Hangya, and Kepecs 2016), it remains unclear how closely implicit and explicit confidence assessment are aligned. Our results reveal two key features regarding the relation between implicit and explicit confidence reports, (i) time investment and explicit reports were strongly related, and (ii), neither confidence report fully predicted the other.

After specific decisions, we requested subjects to assess their confidence using a five-point scale, after their time investment. Subjects’ explicit confidence reports showed signatures of statistical decision: Explicit confidence reports predicted accuracy (single subject example (Figure 6A), *ρ* = 0.98 (Spearman rank correlation), p = 0.005; pooled subjects (Figure 6E), *ρ* = 0.95, p = 0.01), confidence increased with evidence in correct trials (slope > 0, p < 0.001, linear regression, for single subject example and pooled subject data) but decreased in incorrect trials (slope < 0, p < 0.01) (Figure 6B,F), and high-confidence trials showed a steeper psychometric curve than low-confidence trials (Δslope > 0, logistic regression, p < 0.01 (bootstrap) for single subject example and pooled subject data) (Figure 6C,G). Investment efficiency *α* was higher than zero for all subjects (α > 0, p < 0.001 for each subject, bootstrap) and large for most subjects (α > 0.5 in 4/5 subjects) (Figure 6D,H).

We found a strong correlation between time investment and explicit confidence reports in both correct (ρ = 0.53 (Spearman rank correlation), p < 0.001) and error trials (ρ = 0.53, p < 0.001), suggesting a shared confidence computation (Figure 6I,J). For trials in which we obtained both a time investment and explicit confidence report, we could assess whether time investment was correlated with explicit confidence reports beyond their relation to evidence and choice. Time investment in single trials predicted explicit confidence reports even after accounting for the correlation produced by sensory evidence and choice (Figure 6K, stepwise linear regression, p < 0.001).

The same computational process could drive both investment behavior and explicit confidence reports, but we wondered whether this correlation could be explained by subjects inferring their confidence from their own investment behavior, since verbal reports were obtained after investments. We found that explicit confidence reports are not simply a function of time investment, for a fixed time investment, or vice versa. When only considering trials with an explicit confidence report of 3 (the middle of the 5-point scale), time investment was positively correlated with evidence in correct trials (slope > 0, p = 0.05, linear regression) and negatively correlated in error trials (slope < 0, p = 0.02) (Figure 6L). On the other hand, when only considering time investment trials around the median (0.4–0.6 quantile), explicit reports were positively related to evidence in correct trials (slope > 0, p < 0.001, linear regression), while not significantly related to evidence in error trials (p > 0.1) (Figure 6M). These results suggest both confidence-reporting processes were subject to independent ‘metacognitive’ noise. Together, these results suggest that implicit and explicit confidence reports are driven by the same computational process, realizing confidence computations to guide behavior.

### A drift-diffusion algorithm produces near-optimal time investment behavior

To demonstrate the simplicity of optimal time investment strategies, we show that accurately inferring confidence alone suffices—a generic drift-diffusion algorithm can readily transform confidence into optimal time investments. We derived the properties of a drift diffusion model (DDM) to solve the optimal time investment problem (Figure 7A, see Methods).

**Figure 7.**
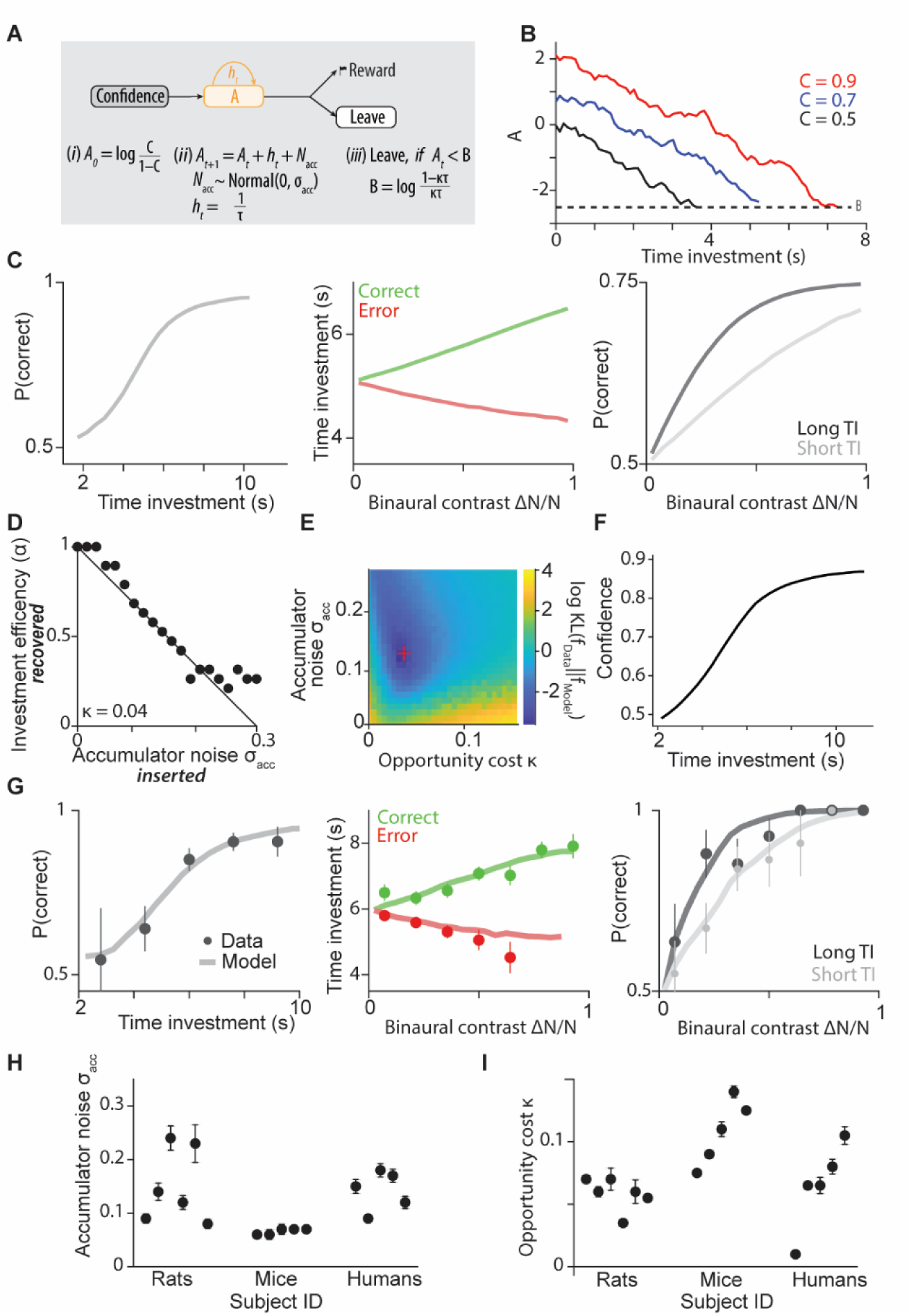
Drift-diffusion algorithm produces near-optimal investment behavior limited by investment efficiency. **(A)** Drift-diffusion model: An accumulator’s initial value A_0_ is proportional to confidence C. During time investment, after each time step Δt, the hazard rate of reward h_t_ is subtracted from the accumulator A_t_, and accumulator noise N_acc_ is added. The agent leaves, if the accumulator’s value falls below a threshold B, which is determined by the opportunity cost of time κ and reward delay distribution parameter τ. **(B**) Progression of the accumulator A for three example decisions with different confidence values. **(C)** Drift-diffusion model produces optimal time investment behavior guided by confidence. **(D)** Investment efficiency captures accumulator noise. **(E)** Model information loss of two-parameter model fit to behavioral data. **(F)** Mapping function from best-fitting behavioral model. **(G)** Predicted time investment behavior from best-fit drift-diffusion model in an example rat. **(H)** Best-fit accumulator noise across single subjects. **(I)** Best-fit opportunity cost across single subjects. Error bars represent SEMs.

The DDM contains an accumulator variable A that starts at A_0_ = log(C/(1-C)), set by the initial confidence C in obtaining a payoff. This represents the log odds ratio of confidence, which linearly accumulates evidence for or against receiving a payoff over time. Using the log odds enables additive integration of the multiplicative effects of confidence on the reward delay distribution that together determine the momentary hazard rate of reward ρ(*t*) (cf. derivation of eq. 7 in Methods). With each passing moment that a reward is not received, the accumulator drifts downwards at a rate proportional to the momentary expected reward rate in correct trials, h_t_. The hazard rate of the reward delay distribution h_t_ corresponds to the probability that a reward will be received at time t in correct trials, given no prior receipt of a reward. Thus, A_t_ will drift more strongly towards the threshold when h_t_ is high, whereas when h_t_ is low, A_t_ will only drift slowly towards the threshold, thus reflecting the decreasing likelihood of payoff over time. In essence, A_t_ tracks the falling likelihood of reward over time. For our task, the reward delay distribution is exponential, and its hazard rate is constant with h_t_ = 1/τ (defined above 0.5 s). When the accumulator A_t_ hits a threshold B, the subject stops its time investment at time t in favor of proceeding to the next trial. The threshold B depends primarily on the opportunity cost of time κ, where a larger opportunity cost implies a larger threshold and therefore subjects leaving earlier. In addition, the threshold depends on properties of the reward delay distribution, here, summarized by the exponential delay parameter τ.

The only source of variability is Gaussian noise, N_acc_, with standard deviation σ_acc_, that is added at each time step to A_t_. For higher confidence level C, the accumulator starts at a higher initial value while drifting at the same rate, therefore leading to longer investment decisions (Figure 7B). As expected, this algorithmic investment model produces optimal time investment (Figure 7C). Deviations from optimal time investment with respect to using confidence to guide investments are captured by the accumulator noise N_acc_. This model does not allow for arbitrary utility functions of time and reward and therefore does not allow us to compute an upper bound on optimality. Instead, we characterized the relationship between the accumulator noise N_acc_ and the investment efficiency α. As expected, higher N_acc_ implied lower investment efficiency α (Figure 7D). We fit two free parameters to behavioral investment data, accumulator noise magnitude σ_acc_ capturing sub-optimality in time investment, and the opportunity cost κ, capturing the overall utility of time and reward (Figure 7E, see Methods). Model predictions quantified the mapping function between time investment and confidence (Figure 7F) and predicted the subjects’ investment behavior (Figure 7G). Across subjects and species, we found variable accumulator noise and opportunity costs (Figure 7HI). For example, mice displayed larger opportunity cost, reflecting their overall shorter time investment behavior (Figure 7I). Together, this investment model algorithmically realizes bounded optimality of time investment.

## Discussion

Here we demonstrate that humans, rats, and mice can use confidence judgments to optimize time investment decisions under uncertainty. By adopting a psychometric decision task with post-decision time investments, we could systematically titrate uncertainty about future payoffs, which allowed us to predict optimal time investment guided by decision confidence. Our computational framework isolates the accurate use of evidence for time allocation but independent of subjective aspects, which we call ‘bounded optimality’, complementing the notion of bounded rationality arising from limited resources (Simon 1997). This comparative approach provides insight into the cognitive architectures that enable rational time investment and metacognitive capacities.

### Bounded optimality across species

Our framework advances the assessment of bounded optimality in three major ways. First, by focusing on the contribution of a specific decision variable, confidence, we quantify optimality independent of assumptions about subjective valuation or utility. Second, the approach isolates a distinct mechanism – the use of confidence signals – complementing the notion of bounded rationality emerging from limited cognitive resources. Third, our experience-based economics task enables cross-species comparison of investment strategies.

It is challenging to determine whether subjects are optimal with respect to their own objective function since, in general, a subject’s utility of reward, effort, and time are unknown (Bach and Dolan 2012; Kable and Glimcher 2009). We addressed this challenge by describing the utility function of time and reward pertaining to confidence using the mapping function. We inferred each subject’s mapping function by only assuming (i) a monotonic relationship between confidence and time investment, and (ii), a stable mapping function across trials and sessions. Even though the mapping function might not be entirely stable across time, variability in the mapping function would lead to a lower apparent investment efficiency since confidence noise would partially originate from this variable. Therefore, our investment efficiency estimate provides a lower bound to the subject’s actual use of decision confidence to guide investment decisions.

Further, each subject’s ability to make predictions about rewarded choices was limited by their sensitivity to auditory discriminations (‘perceptual noise’), producing variability in investment behavior. Our optimal model predictions assume that agents fully leverage sensory uncertainty by computing confidence to guide time investment. Any further limitations of the agent’s cognitive capacities such as time estimation or the metacognitive use of confidence will lead to sub-optimal performance. Hence, our bounded optimality model provides an upper limit of investment performance with respect to decision confidence. We do not know if time investment is also optimized with respect to other sources of uncertainty such as known risks, uncertainty in subjective value.

The cross-species similarities in confidence-based investments are striking given the radically different evolutionary histories. Rats and mice, like humans, appear adept at using metacognitive signals to optimize economic decisions. Despite cognitive limitations, all three species achieved near-optimal performance by investing proportional to their decision confidence. The capacity to fine-tune investments based on internal confidence assessments may thus be an evolutionarily ancient competence. Since we only investigated one type of investment behavior, namely investing time after committing to a choice, we cannot generalize across other types of investments, such as money or effort. However, the same computational framework of statistical confidence can be used to study other sources of uncertainty as well as other forms of investment behavior. Our results thus provide a framework to study investment behaviors across species.

### Optimal time allocation in decision-making

Time allocation is a fundamental problem in decision-making and has been studied primarily in two domains. First, as a reflection of a speed-accuracy trade-off processes when deciding between choice alternatives. Second, in optimal foraging theory when deciding how much time to spend at resource patches. In either case, and our time investment task, the core challenge is that expending more time can increase the chance of receiving a payoff, but at the cost of missing out on other opportunities, i.e., paying an opportunity cost (Becker 1965; Hausfeld and Resnjanskij 2018; Rustichini 2009).

In decision-making, the speed-accuracy tradeoff posits that taking more time to make a decision leads to more accurate outcomes (Wickelgren 1977). Drift-diffusion models implement optimal decisions in the sense that decision time is minimized to achieve a certain accuracy level, and choosing an appropriate decision threshold maximizes reward rate (Bogacz et al. 2006; Ratcliff 1978). Here, more difficult decisions always take more time. However, it is sub-optimal to spend more time deliberating when decision alternatives have similar expected value and either choice leads to similar payoffs, complicating the relation between time allocation and reward maximization in decision-making (Oud et al. 2016; Pirrone, Stafford, and Marshall 2014). In contrast, optimal time allocation in our post-decision investment task directly constructs a tradeoff between the opportunity cost of investing and the expected payoff, which is the probability of being rewarded (i.e., decision confidence). Variation in time investment after each decision can therefore be assessed through the lens of variable decision confidence.

Time investment, or how much time you spend in pursuit of a future gain, has been studied as the problem of optimal time allocation in animal foraging (Stephens and Krebs 1986; Kacelnik 1997). A major theorical result, the marginal value theorem, posits that optimal time allocation in depleting food patches should be in proportion to the average reward rate in the environment, which directly corresponds to the opportunity cost of waiting (Charnov 1976; Constantino and Daw 2015). While many studies found empirical evidence for the behavioral strategies aligned with the marginal value theorem, quantitative deviations from the theory arise particularly in time-limited foragers, or when time perception is distorted (Hills and Adler 2002; Nonacs 2001; Wajnberg et al. 2006). Deviations from optimality predictions become difficult to interpret, since the subjective cost of waiting and the discount rate of future rewards are generally unknown (Hayden 2016). Even if the subjective cost of waiting can be evaluated in economic decision tasks in humans (Cruz Rambaud, Ortiz Fernández, and Parra Oller 2023; Ventre and Martino 2022) or animals (Sweis et al. 2018; Wikenheiser, Stephens, and Redish 2013), lack of normative decision models that take into account trial-by-trial variations in the subject’s uncertainty to receive a reward make it challenging to interpret sub-optimal behavior (Ott et al. 2022). The strength of our post-decision investment task is that it allows quantification of the subjective uncertainty (decision confidence) trial-by-trial, and therefore we can assess to what degree subjects appropriately used their sense of confidence to adjust graded time investment decisions. In addition, since we were able to non-parametrically estimate the mapping function between confidence and time investment, we circumvent the need to make strong assumptions about the shape of the subjects’ utility function of reward, effort, and time. Future studies that are designed to independently measure a subject’s utility of time and reward could establish to what degree time investment decisions maximize the subject’s general utility.

### Investment decisions and temporal discounting in behavioral economics

In behavioral economics, investment decisions have been typically studied using explicitly described, hypothetical scenarios and decisions made by responding to a survey or questionnaire. In many cases, these studies revealed that participants fail to make rational investment decisions, distorting the risks and payoffs of future gains (Camerer 1999; Hirshleifer 2015; Tversky and Kahneman 1981). Such risk attitudes are considered general, temporally stable traits (Dohmen et al. 2011; Frey et al. 2017). How can we reconcile the findings about irrational economic decisions with our claim about near-optimal investment decisions? First, an important difference in our study is that subjects enacted investment decisions, experienced their consequences, and could learn from previous decisions. This experiential component, which was further enhanced by gamifying human tasks, might facilitate rational investment decisions and contribute to an description-experience gap between explicitly described decision scenarios and behavioral tasks with experiential structures (Basile, Fabien, and Stefano 2020; Hertwig and Erev 2009; Wulff, Mergenthaler-Canseco, and Hertwig 2018). Second, we believe that expending an actual cost – limited time – contributes to establishing an incentive structure that facilitates the use of time as a valuable resource, such as money (Festjens et al. 2015; Zauberman and Lynch 2005; Etkin, Evangelidis, and Aaker 2015), which might lead to more accurate confidence representations (Lejarraga and Lejarraga 2020; Kaufmann, Weber, and Haisley 2013) and can be leveraged across species. Finally, our behavioral task to study investment decisions allows for psychometric control and model-based assessment of the uncertainty that contributes to investment decisions. It is important to note that we claim near-optimal behavioral with respect to using this uncertainty to adjust behavior – bounded optimality –, rather than assessing potential distortions in representing value or risk as in many studies in behavioral economics (Camerer 1999; Hirshleifer 2015; Tversky and Kahneman 1981). Similar in spirit, our notion differs from ‘bounded rationality’ (Simon 1997; Sent 2018) in that apparent irrational behavior can be optimal in light of highly subjective utility function but with respect to subjective sources of evidence. Two main sources of subjective utility functions are (1) risk distortions as in prospect theory and observed in rats (Constantinople, Piet, and Brody 2019) and non-human primates (Stauffer et al. 2015; Yang, Li, and Stuphorn 2022), besides humans, and (2) temporal discounting as observed across species (Hayden 2016; Vanderveldt, Oliveira, and Green 2016; Cruz Rambaud, Ortiz Fernández, and Parra Oller 2023). In our study, such distortions in utility functions as well as temporal discounting are captured by our mapping function. For example, explicit confidence reports showed an approximate linear relationship with accuracy, while large time investment predicted only small increases in accuracy, which could reflect additional factors that determine time investment such as the motivation to wait or uncertainty in time estimation. The shape of the mapping function captures the subjects’ time preferences, i.e., whether they are impulsive or impatient as a function of their confidence. Time preferences can be adaptive, depending on uncertainty, risk, and opportunity costs (Boon-Falleur, Baumard, and André 2021; Mell, Baumard, and André 2021; Fenneman, Frankenhuis, and Todd 2022). Our approach allows for systematic quantification of time preferences across species or individuals.

### Time investment as a behavioral report of confidence

Because time investment decisions were guided by decision confidence, they served as a behavioral report of confidence across humans, rats, and mice. These results build on previous findings that rat time investment followed the signatures of statistical decision confidence in auditory and olfactory discriminations (Lak et al. 2014; Masset et al. 2020; Ott, Masset, and Kepecs 2019; Stolyarova et al. 2019; Joo et al. 2021), and mouse time investment reflected statistical confidence in an auditory detection task (Schmack et al. 2021). We have also previously found that human explicit confidence reports reflected statistical decision confidence (Sanders, Hangya, and Kepecs 2016). Here, we were able to directly compare ‘implicit’ confidence reports (i.e., time investment) and ‘explicit’ confidence reports (i.e., verbal reports) in single trials. The tight correlation between explicit and implicit reports suggests that the same core computation, statistical decision confidence, underlies both reports. Thus, time investment serves as a robust behavioral readout of confidence across species.

Confidence has been studied through several behavioral studies in humans and other animals. Most behavioral tasks use difficult binary choices (e.g., sensory discrimination) that provide a third option, for example to opt out, to obtain a small but guaranteed payoff (Foote and Crystal 2007; Grimaldi et al. 2018; Hampton 2001; Komura et al. 2013; Middlebrooks and Sommer 2011; 2012; Shields, Smith, and Washburn 1997). We believe that our post-decision investment task affords several advantages (Kepecs and Mainen 2012). First, we obtain a decision and confidence report in the same trial, allowing us to directly assess to what degree confidence predicts accuracy. Second, we obtain a graded measure of confidence, allowing us quantitatively relate time investment with task variables such accuracy, choice difficulty, and choice correctness. Finally, psychometric control of the subject’s choice allows us to probe confidence along a continuous range of evidence strength and choice. Together, this approach enables us to leverage a statistical model of confidence (Hangya, Sanders, and Kepecs 2016) and quantitatively relate evidence, choice, confidence, and time investment.

We assessed investment efficiency by quantifying deviations from optimal time investment predictions. This approach is related to determining the ‘metacognitive efficiency’ of confidence ratings in human subjects, asking how well confidence ratings predict accuracy (Fleming and Dolan 2014; Fleming and Lau 2014). Similarly, we assessed how well time investment predicted accuracy. Inherently, monitoring accuracy is limited by the subject’s sensitivity to discriminating sensory stimuli. By using a statistical model of confidence (Hangya, Sanders, and Kepecs 2016), we could predict the distribution of expected confidence values for each stimulus and choice given the subject’s sensory sensitivity. Our model thus allows for comparing investment efficiency across subjects with unequal sensory sensitivity, similar to other model-based approach such as meta d’ (Maniscalco and Lau 2012) or measuring the difference in psychometric slopes between low and high confidence trials (De Martino et al. 2013), an analysis we included as part of our key behavioral signatures. Our generative model allows for single-trial predictions of optimal time investment distributions, or, more generally, confidence reports fully informed by statistical decision confidence. This avoids the necessity to rely on binary comparisons such as sensitivity between two stimuli or sensitivity between low and high confidence reports required for many other measures, which are therefore prone to high variability (Bor et al. 2018; Fleming and Lau 2014). Similar to our approach, recent generative models fit metacognitive noise together with other behavioral parameters (Boundy-Singer, Ziemba, and Goris 2023; Guggenmos 2022). In contrast, here, we first estimated all behavioral parameters not related to metacognitive noise to subsequently estimate an upper bound for metacognitive noise, or bounded optimality. Our approach allows for explicitly fitting generative noise models and could thus enable future studies to delineate sources of metacognitive noise.

### Drift diffusion investment algorithm for bounded optimality

Algorithmic models can link normative models with their candidate implementation in neural circuits (Love 2015; Marr 1982). We therefore investigated whether drift diffusion models (DDMs) – a large class of models commonly used in decision-making (Ratcliff 1978; Bogacz et al. 2006; Gold and Shadlen 2007; Brunton, Botvinick, and Brody 2013) – can implement bounded-optimal time investment. Motivated by the fact that decision confidence directly follow from the subjects’ percept of the sensory evidence and choice (Hangya, Sanders, and Kepecs 2016; Lak et al. 2014), we found that generic DDMs set to initial values that only depend on decision confidence can implement optimal time investment. The properties of the reward delay distribution and the opportunity cost of time were captured by the drift rate (equal to the hazard rate of the reward delay distribution) and the threshold. It is thus conceivable, that the neural decision confidence signals present at the time of choice, when investment begins, form the starting point of dynamic ramping activity during waiting (Ott, Masset, and Kepecs 2019; Masset et al. 2020). Our algorithmic model would further predict neural signatures of the accumulator, akin to a ramp-to-threshold mechanism (Shadlen et al. 2001; Gold and Shadlen 2007; Pinto, Tank, and Brody 2022; Koay et al. 2022), and neural signatures of the reward delay distribution. Such neural data would further inform and constrain algorithmic models that implement bounded optimality of time investment behavior across species.

In conclusion, this study establishes a new approach to comparatively probe the neurocomputational bases of economic rationality. Despite cognitive limitations, simple statistical representations, like confidence, may enable mammals to robustly optimize decisions and approach bounded optimality in key domains like investing time. Quantitative models clarifying the mechanisms and evolutionary origins of such competencies will offer insights into both animal and human cognition.

## Acknowledgments

This work was supported by the German Research Foundation (DFG) OT562/2-1 and the European Research Council (ERC) Starting Grant TIMEVALUE to TO. AK was supported by NIH R01DA038209 and R01MH097061.

## Author contributions

TO and AK conceived, designed and interpreted the study. TO performed experiments (rat data), analyzed the data, and wrote the manuscript. JIS, MB, PM performed experiments (JIS: human data, MB: mouse data, PM: rat data) and contributed to experiment design and conceptualization. AK supervised the project and edited the manuscript. All authors provided comments on the manuscript.

## Methods

### Human experiments

#### Subjects

Five human subjects performed 27,212 trials across 74 behavioral sessions, recruited from the general population of Cold Spring Harbor Laboratory. Subjects ranged from 23 to 30 years of age and reported normal or correct-to-normal vision and had no known of hearing impairments. Each participant provided written informed consent prior to participation, with all procedures approved by the Cold Spring Harbor Laboratory Institution Review Board.

#### Apparatus

A single apparatus was available to the subjects in a dark room with 24-hour access using their employment card and subjects could choose when to complete one session. Instrument control and data acquisition were accomplished in MATLAB (R2011a) using the statistics and psychophysics toolboxes. Computer peripherals included a USB joystick (ST290, Saitek), a standard USB keyboard (Dell) and an LCD monitor (G245HQ, Acer) positioned at eye level, 70 cm in front of the subject. Subjects initiated a trial and thus stimulus delivery by squeezing the joystick trigger and reported their choice (left or right) on the USB keyboard arrow keys. For explicit confidence reports, custom-printed adhesive labels were placed over the keypad’s numerical keypad to indicate confidence levels “1” to “5” on a 5-point scale, presented vertically to capture the intuition about low and high confidence levels and to discourage confidence-side associations. An additional button labelled “Err” was provided to allow the subject to indicate a choice believed to be wrong. Trials with “Err” responses were pooled with confidence report of 1.

A real-time click generator and response capture device (PulsePal, Sanworks, (Sanders and Kepecs 2014)) delivered the auditory stimulus binaurally to headphones worn by the subject (HD-280, Sennheiser). Pulse Pal’s firmware was modified to record the subjects’ joystick and keyboard press and release events with high temporal precision (100 µs resolution). A pair of computer speakers was connected to the computer’s mainboard sound port and positioned on either side of the subject beneath the monitor to play back feedback sounds.

To encourage long term participation, a custom appetitive reward system was installed above the computer. Three servo motors (HSR125-CR, Hitec) controlled by a USB servo controller (Micro Maestro, Pololu) motorized a confection dispenser (Mini Bean Machine, Jelly Belly USA). Plastic pipes delivered the dispensed items to a plastic dish on the subject’s left side at arm level. The appetitive reward system was only used in the break periods between blocks of trials (see below).

Data were automatically labeled by unique subject ID codes to ensure anonymity. A set of keys relating subject ID codes to subject identities was encrypted and stored on a removable drive.

#### Subject instructions

Subjects verbally agreed to complete at least eight sessions, and at least two sessions per week in consecutive weeks. Use of the apparatus was demonstrated prior to the first session and an instruction sheet explaining button and keypad usage was posted in the test room.

Subjects were paid $ 0.10 for each correct response and were informed that random guessing would earn them $7 per hour on average if they responded continuously, and that accurate responses could earn them more than $20 per hour. Subjects were periodically paid in cash, when they requested a payout, or when their balance exceeded $200. Such large incentives were necessary to encourage subjects to optimally use time within trials. Subjects were encouraged to use the entire confidence scale and were advised that they may need to ‘remap’ their initial range of experienced confidence to span the 5-point confidence reporting scale. Subjects were not given explicit speed or accuracy instructions. Rather, in the interest of similarity to the rodent task, subjects were instructed to earn as much payment as they could in the time allotted. The first two sessions used for training and to adjust discriminability levels to subject performance and were not included in subsequent analyses. Following training sessions only, subjects were given written feedback about side bias, scale usage and performance. Subjects typically learned instructions immediately, reached peak performance within two sessions, and did not substantially improve their performance in subsequent test sessions.

#### Time investment task in humans

We designed a perceptual 2-alternative forced choice task complemented with a post-decision temporal wager to record both a choice and a time investment in a single trial. Subjects started a trial by pressing the main joystick button. Binaural auditory stimuli were delivered after random delay (exponential random delay with time constant *τ* = 1.5 s) via headphones. In each ear, subjects listened to a series of Poisson-distributed clicks with different mean underlying rates. Subjects had to determine the side (left or right) with the higher underlying click rate and indicate their choice by pressing a button on the response device. Pressing the response button instantly terminated the click train. For each trial, we randomly chose a delta click rate between left and right from six levels between 0 and 100. The sum of the left and right click rate was kept constant at 100. Click times were drawn randomly for each trial using a Poisson distribution and delivered with the PulsePal (Sanworks, NY) pulse generator to the headphones. The six possible underlying click rates were manually adjusted symmetrically around 50 clicks/s such that the stimulus strengths for each subject after the first two sessions achieved approximate performance levels of 60 %, 75 % and 95 %. In addition, we added 10 % of neutral evidence trials with an underlying click rate of 50 clicks/s for both left and right, which were randomly rewarded. Since reaction times varied across trials, we re-calculated the per-trial click rate based on the number of clicks presented to the subject during stimulus playback.

To measure time investment behavior, subjects had to keep pressing the response button (left or right) after making a choice for a random amount of time to receive a money reward for correct choices, which was added to a virtual account. Random delays were drawn from a truncated exponential distribution (minimum, 0.5 s; maximum, 8 s; time constant *τ* = 1.5 s). We chose the exponential distribution to maintain a relatively constant reward expectation, i.e. a flat hazard rate. For correct trials, after the random delay passed, a visual feedback and feedback tone indicated a correct response and the cumulative earned amount of money earned during the session was displayed on the screen before the next trial was initiated. There was no feedback on error trials and in a subset of correct trials (probe trials, 10 %, randomly interleaved). In these trials (error and probe trials), subjects kept pressing the response button for a self-determined amount of time, thus investing time in their decision. The time from pressing until releasing the response button thus constitutes a time investment. In all probe trials and 30% of error trials, subjects were prompted to explicitly report their confidence using a 5-point scale on the keypad. In a pilot study, most human subjects showed poor time investment behavior, i.e., time investment poorly calibrated to decision confidence. In contrast to rodent studies, the incentive structure in human psychophysical tasks is less clear. In particular, we sought to overcome (i) difficulty in task engagement given that hundreds to thousands of trials were necessary for model-based analysis, and (ii) ambiguous incentive structures with unlimited resources and decorrelation between performance and reward. We therefore designed a video game interface for psychophysics, previously suggested to encourage task participation (Abramov et al. 1984; Soderquist and Shilling 1992). Gamification created clear incentive structures by limiting the time to collect reward to brief 5-minute blocks (max. 15 blocks/session). Therefore, optimizing time allocation will yield more money. In addition, subjects reported improved task engagement, and higher rates of returning for additional sessions. In our “Plutonium miner” game the subjects’ goal was mine valuable Plutonium by locating radiation sources using a Geiger counter (since the Poisson click streams resembled clicks on a Geiger counter). Subjects controlled an avatar on a game map with several “mining sites” each representing a block of trials. To access each site, subjects needed to purchase access at a “land office” building at the center of the map, where subjects were reminded to earn as much reward as possible at each mining site. On entering a site, subjects committed to a five-minute block of trials starting immediately. After initiating a trial, a signpost in a static field of dots was shown. The dot field was randomly generated on each trial. During stimulus presentation, the signpost was removed, and the subject’s avatar was displayed within the dot field. No visual changes occurred during stimulus presentation and visual displays did not predict sensory evidence. After making a choice and during waiting, the avatar was facing the chosen side while the random dot field was moving in the opposite direction to provide the visual impression of the avatar moving towards the chosen side. If rewarded, the avatar jumped repeatedly, and the reward amount was shown flashing above its head. For each correct trial, the subject earned 200 units of virtual currency, and the cost to purchase access to the next block of trials was determined as ¼ of the subject’s total profit from the previous block. To promote engagement within a session, subjects could spend virtual currency to produce candy at virtual factories (M&Ms, Skittles or Reeses Pieces). Subjects were notified in subsequent inter-block intervals if candy production was completed, which was then delivered to the subject by the appetitive reward apparatus.

Overall, gamification produced stable choice and time investment behavior. Mean performance, explicit confidence and time investment across blocks were not correlated with the currency spent in the previous block interval. Thus, between-block game elements or behavior did not impact time choice or investment behavior within blocks.

#### Data analysis

We collected 27,212 trials from five humans across 74 behavioral sessions (subjects performed between 5 and 24 sessions each, number of trials within a session averaged 395). We did not analyze the first 50 trials of each session and trials with a time investment smaller than 1 s were excluded. For auditory stimuli, we computed the binaural contrast *x = (N_Left_*–*N_Right_)/ (N_Left_+N_Right_)* using the number of clicks presented in each trial. We fitted auditory choices using a cumulative Gaussian distribution. Error bars represent 95 %-confidence intervals across trials unless otherwise noted. For the calibration and conditioned psychometric curves (Figures 3-5A,C,E,G), we used all correct probe trials and randomly re-sampled from error trials to match the subject’s average performance. For the vevaiometric curve (Figures 3-5B,F), we included all error trials. The calibration curve was statistically evaluated using Spearman rank correlation by using 1 s time investment bins or nominal explicit reports. The vevaiometric curve was evaluated by linear regression of time investment (or explicit reports) against absolute evidence |*x*| separately for correct and error trials (*t*-statistic). The conditioned psychometric curves were evaluated by fitting a 3-parameter logistic regression for choice against absolute evidence (slope, bias, lapse) separately for high-confidence and low-confidence trials (split by median time investment) and statistical significance of the difference in slopes was estimated using bootstrap simulations (N = 999). For assessing whether time investment and explicit confidence reports were related beyond evidence and choice, we used stepwise linear regression. First, we calculated the residuals of a linear regression models of time investment against absolute evidence |*x*| separately for correct and error trials, and regressed explicit confidence reports against residual time investment, assessing whether this relation was statistically significant and positive (*t*-statistic). Data analysis was performed using custom software (MATLAB).

### Animal experiments

#### Subjects

We used 6 adult (> 12 weeks) male rats (Long Evans, Taconic) housed in pairs and maintained on a reverse 12 hr light/dark cycle and 5 adult (> 10 weeks) (3 female, 2 male) mice (C57BL/6JAX, Jackson Laboratory). Rats and mice had ad libitum access to food and were under a liquid restriction schedule with daily monitoring of water intake to maintain a body weight of at least 85 % of free-drinking weight. Animals obtained water during daily training sessions and received ad libitum water on weekends and as needed. All procedures involving animals were carried out in accordance with National Institute of Health standards and were approved by the Cold Spring Harbor Laboratory Institutional Animal Care and Use Committee.

#### Apparatus

Behavioral procedures were conducted similarly as previously described (Masset et al. 2020). A rectangular behavioral box contained three ports aligned on one wall equipped with LEDs, infrared photodiodes, and phototransistors. Interruption of the infrared photo beam was used to determine port entries and port exits. The two side ports (choice ports) were additionally equipped with valve-controlled water spouts for reward delivery. Auditory stimuli were delivered using the pulse generator PulsePal (Sanworks, NY) and two speakers placed symmetrically outside of the behavioral box at the left and right panel, aligned to the animal’s head. Water valves, LEDs and phototransistors were controlled by the behavioral measurement system Bpod (Sanworks, NY) and custom software (MATLAB).

#### Time investment task in rodents

In a time investment task analogous to human experiments, we designed a perceptual 2-alternative forced choice task complemented with a post-decision temporal wager to record both a choice and a time investment in a single trial from rats and mice (Masset et al. 2020). The first part of the trial was constructed as a two-alternative forced choice task. Animals self-initiated each trial by entering the central stimulus port. After a random delay of 0.2–0.4s, an auditory stimulus was presented. Animals had to determine the side with the higher number of clicks in binaural streams of clicks. Auditory stimuli were random Poisson-distributed click trains played binaurally at the two speakers placed outside of the behavioral box for a fixed time of 0.35 s. For each subject, we chose a maximum click rate click_max_ according to the performance of the animal, typically 50 clicks/s (full range 40–100 clicks/s). This maximum click rate was fixed for each animal. For each trial, we randomly chose a delta click rate between left and right from a uniform distribution between 0 and click_max_. The sum of the left and right click rate was kept constant at click_max_. Click times were drawn randomly for each trial using a Poisson distribution. For a subset of mice (M1, M2), we instead used 6 fixed click rates distributed symmetrically around click_max_/2, which were manually chosen during training to match average accuracy of approximately 60 %, 70 %, and >90 %. Stimuli were delivered with the PulsePal (Sanworks) pulse generator to the speakers placed left and right at the behavioral chamber. Animals indicated their choice by exiting the stimulus port and entering one of two choice ports (left or right) with a maximum response time of 3 s after leaving the stimulus port. Choices were rewarded according to the higher number of clicks presented between the left and right click train (equal number of clicks were randomly rewarded). Exiting the stimulus port during the pre-stimulus delay or during the stimulus time (first 0.35 s) were followed by a white noise and a time out of 3–7 s. In rats, we randomly interleaved trial with olfactory stimuli as previously described (Masset et al. 2020). Olfactory trials were excluded from all analyses in the present study.

To assess the rats’ and mice’ degree of confidence, we asked animals to place a temporal wager on their decision by investing a self-determined amount of time after committing to a choice. For correct choices, we withheld reward delivery for a random time, drawn from a truncated exponential distribution (rats: minimum = 0.5 s, maximum = 8 s, time constant *τ* = 1.5 s; mice: minimum = 0.5 s; maximum = 5 s; time constant *τ* = 1 s). The time distribution for mice was chosen to be shorter because mice tended to show greater difficulty in waiting. We chose the exponential distribution to maintain a relatively constant reward expectation, i.e., a flat hazard rate. Rats and mice had to keep poking the choice ports during the entire time period. On a subset of trials (probe trials; rats 10 %, mice 5 %), we withheld reward entirely. In addition, there was no feedback on error trials. Consequently, animals decided to leave the choice after a variable amount of time to initiate the next trial. To avoid false detections of leaving decisions animals had a grace period of 0.3 s in which re-entry into the side port was not considered a leaving decision. After a leaving decision was complete, a tone indicated to the animal that the trial finished. This provided us with a continuous time investment as well as a binary choice on each non-rewarded trial (probe trials and error trials). Time investment served as an index of the rats’ confidence estimate in their decision (see below).

#### Training

We first started training naïve rats or mice to enter the central stimulus port and subsequently collect a water reward at either of the two choice ports (left or right). Animals learned to enter the stimulus port for at least 0.7 s. Subsequently we introduced auditory stimuli, starting with stimuli of high discriminability. After performance levels reached above 90 % we introduced stimuli with decreasing discriminability, i.e. increasing difficulty, to sample the full range of the animal’s choice behavior. During this early training stage, all correct choices were rewarded and incorrect choices were followed by a feedback of white noise and a time out (3–7 s). After stable performance levels (lapse rates <5–10 %) we removed feedback for error trials and introduced randomly delayed rewards gradually, starting from a maximum of 2 s to a maximum of 8 s over the course of several sessions. We never differentially reinforced leaving behavior at the choice port. Finally, we introduced probe trials. Overall training took typically between 40–60 training sessions over the course of 8–12 weeks. Each session lasted about 2-3 hours, in which the subjects typically completed 500 trials.

#### Data analysis

For rats, we collected 32,391 trials from six rats across 98 behavioral sessions (rats performed between 17 and 24 sessions each, number of trials within a session averaged 563). For mice, we collected 94,991 trials from five mice across 217 behavioral sessions (mice performed between 10 and 77 sessions each, number of trials within a session averaged 448). We did not analyze the first 50 trials of each session and trials with a time investment smaller than 2 s (rats) or 1 s (mice). For auditory stimuli, we computed the binaural contrast *x = (N_Left_*–*N_Right_)/ (N_Left_+N_Right_)* using the number of clicks presented in each trial. We fitted auditory choices using a cumulative Gaussian distribution. Error bars represent 95 %-confidence intervals across trials unless otherwise noted. For the calibration and conditioned psychometric curves (Figures 4-5A,C,E,G), we restricted our analysis to probe trials to keep the average accuracy equivalent to the rest of the trials. For the vevaiometric curve (Figurs 4-5B,F), we included all error trials. The calibration curve, vevaiometric curve, and conditioned psychometric curves were evaluated as for the human data (see above). Data analysis was performed using custom software (MATLAB).

### Confidence-guided time investment model

#### Rationale

We developed a confidence-guided time investment model to (*i*) assess whether human or rodent subjects invested time according to their degree of confidence, and (*ii*) quantify deviations from optimal time investment. Crucially, optimal time investment in our post-decision time investment task is guided by each trial’s subjective confidence estimate (eq. (1) – (6)).

Our model is structured into 4 steps. First, since we systematically manipulated the evidence for a choice across trials, we can infer the subject’s perceptual noise (*Step 1*) and subsequently estimate the subject’s confidence in each trial (*Step 2*). We can then convert confidence into an investment time using a mapping function *T* = *m*(*C*) (*Step 3*). Finally, we can fit a single noise parameter, investment efficiency, to quantify deviations from optimal time investment (*Step 4*).

#### Step 1: Infer perceptual noise σ

To infer each subject’s perceptual noise σ, we assume that a percept *x̂* is a noisy representation of the true evidence strength *x* in a given trial (Gaussian noise with standard deviation σ), and that the choice is based on the percept alone (cf. Figure 2A). For each subject, we fitted the subject’s psychometric curve using a truncated Gaussian cumulative distribution with parameters σ (perceptual noise) and *m* (bias), using least-squares model fitting, and defining evidence strength as the binaural contrast *x = (N_Left_-N_Right_)/ (N_Left_+N_Right_)* using the number of clicks presented in each trial.

#### Step 2: Compute model confidence per trial

We used a statistical definition of confidence to compute model confidence for each simulated trial. Confidence was defined as the probability of being correct given the subjective level of evidence (percept *x̂*) and choice, P(correct|*x̂*,choice). We generated 1,000,000 model samples using the same evidence strength *x* as used in behavioral sessions. We then model ‘percepts’ *x̂* for each sample by drawing from a Gaussian distribution with mean *x* and standard deviation σ. For each percept *x̂* and choice, we then calculated its associated confidence *C* = P(correct|*x̂*,choice), binning *x̂* into 500 equally sized bins and defining an interpolated ‘belief function’ *x̂* → *C*.

#### Step 3: Estimate mapping function m(C)

What is the mapping function between confidence C and time investment *T*, i.e., *T* = *m*(*C*)? According to eq. (6), optimal time investment *T_opt_* is given by

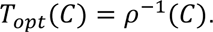

Here, ρ(*t*) is the hazard rate of reward per unit time, i.e., the time-varying probability that reward arrives in the next time interval from *t* to d*t* after waiting time *t* without receiving a reward. Note that ρ(*t*) has a theoretical solution (Lak et al. 2014), which depends on (1) the subject’s confidence in each trial, and (2) the experimenter-defined reward exponential distribution P_rew_(*t*) = *e*^(-t/*τ*)/*τ*^ (*τ* is the temporal decay), i.e.,

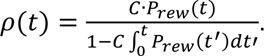

Therefore, *ρ*^−1^(*C*) is given by

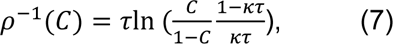

where κ is the opportunity cost per unit time, treated as stable (Lak et al. 2014). Note that we did not change the reward distribution and observed stable investment behavior across many sessions.

Even though, according to (7), the mapping between confidence and optimal time investment has a theoretical solution, we expect the subjects’ subjective mapping function to substantially deviate from this solution. First, scalar timing uncertainty will distort the subjects’ representation of the hazard rate. Second, we cannot be sure with which precision subjects learned the shape of the reward distribution P_rew_(*t*). Thirdly, we do not know the investor’s utility of reward or effort in waiting. Finally, we do not know the interaction between time and utility, i.e., the investor’s value of time. For these reasons, we sought to empirically infer the mapping function between confidence and time investment, which allows us to isolate the contribution of decision confidence to time investment behavior, expressed by

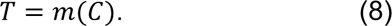

By assuming that *m*(*C*) is a monotonically increasing function, i.e., higher confidence implies higher time investment, we can empirically infer *m*(*C*) based on the subjects’ overall time investment distribution *f_T_*(*T*). Since our model predicts a confidence distribution *f_C_*(*C*), we can consider the transformation *T* = *m*(*C*) as a change of variables. Thus, *m*(*C*) is given by

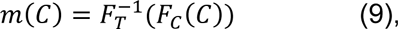

where F*_C_* and F*_T_* are the cumulative distribution functions of model confidence C and observed time investment T, respectively.

To empirically determine *m*(*C*) we first estimating *F_T_*(*T*) using a kernel density estimator (normal kernel function) of the observed time investment distribution (probe trials only) evaluated at 100 equally spaced time investment points and subsequently calculated quantiles of the model confidence distribution *f_C_*(*C*) at F*_T_*(*T*). This yielded the inverse mapping function *m*^−1^(*T*) from *T* → *C*, numerically defined at 100 points. Since we assumed monotonicity, could then converted each model confidence into time investment by defining an interpolated mapping function *m*(*C*) from *C* → *T*.

Together, this model yields an optimal time investment for each trial. Since we do not know the subjects’ utility function of reward and time, note that, here, optimality refers to the statistically optimal use of evidence available to the subject. The strength of our approach is to isolate the statistical optimal use of evidence to time investment behavior, which is required for a reward-maximizing decision strategy.

We constructed 95 %-confidence intervals for *m*(*C*) using bootstrapping by randomly sampling trials with replacement 999 times.

#### Step 4: Fit investment efficiency α

Finally, we quantified deviations from optimal time investment by fitting a single noise parameter, investment efficiency α. Our model yields a distribution of predicted investment values for each evidence and choice combination since we do not have access the subject’s graded percept in a single trial. Nevertheless, this allows computing the likelihood of observed time investment under our model, and therefore using maximum-likelihood techniques to fit a single investment efficiency parameter α and thereby assess deviations from optimality.

Investment efficiency α is defined as the fraction of trials with accurate confidence, while (1–α) is defined as the fraction of trials with random confidence, drawn from the overall model confidence distribution. For the set of model confidence values *C*, a randomly selected subset *K* = (1–α)*N* (*N* = 1,000,000 trial simulations) was replaced by *C_k_* = *C*_shuffle,*k*_ (*k* ∈ *K*) with *C*_shuffle_ representing the set of shuffled confidence values. This corresponds to the fraction of trials, in which the subjects ‘forgot’ their degree of confidence. Note that α provides a convenient index for investment efficiency between 0 and 1 while retaining confidence and time investment distributions (i.e., m(C)), where 0 corresponds to random time investment not related to confidence (and not showing any key signatures of confidence, dashed line in Figure 2F-H) and 1 corresponds to optimal time investment fully informed by confidence.

For each time investment trial *k* (probe trials and incorrect trials), we observed choice *d_k_* and time investment *T_k_* for evidence *x_k_*. We calculated the model likelihood of observing time investment *T_k_* given by *L*_*k*_(*α*) = *P*(*T*_*k*_ |*x*_*k*_, *d*_*k*_, *α*), by using Monte-Carlo simulations described in steps 1-3 to compute model choice *d*, confidence *C*, and time investment *T* = *m*(*C*) distributions for evidence *x* (6 equally sized bins) and time investment *T* (10 equally sized bins). We then determined α_best_ by maximizing the likelihood function 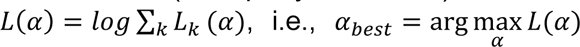, by sweeping across 100 equally spaced investment efficiency values α between 0 and 1.

We constructed 95%-confidence intervals for *α*_*best*_ using bootstrapping by randomly sampling time investment trials (with replacement) 999 times. This also yielded p-values for α > 0. We assessed the model’s capacity to recover optimal time investment (α = 1) by randomly subsampling model trials equal to the number of observed trials 1000 times and observing the resulting best-fit α distribution (gray area in Figures 3,4,5 panel D). Model recovery demonstrated that we could recover the true α by constructing simulated data with infused investment efficiency α between 0 and 1 (Figure 2I). In addition, we tested our model’s capacity to recover other forms of noise. We simulated data infused with three different noise models and used our investment efficiency model to fit α. Investment efficiency α was able to capture all three different noise models. Noise models were (i) a ‘mean-noise’ model with parameter β, in which C_β_ = C·β + C_shuffle_·(1–β) (0 < β < 1), where C_shuffle_ represent randomly shuffled confidence, (ii) ‘Gaussian-noise’ model with parameter γ, in which C_γ_ = C + 𝒩(0, σ = γ) (0 < γ < 4), and (iii) ‘scalar timing’ with parameter Φ, in which C_Φ_ = C + 𝒩(0, Φ·C) (0 < Φ < 0.8).

### Drift diffusion model

To derive an algorithmic investment model that dynamically unfolds across time and can implement optimal investment behavior, we considered drift diffusion models (DDMs) with a single particle. We considered the class of DDMs comprising an accumulator A_t_ and threshold B described by

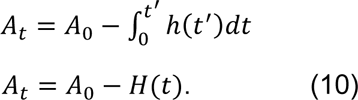

Here, h(*t*) is the hazard rate of the reward delay distribution, i.e., the probability of receiving a reward at time *t* after waiting time *t* in a rewarded trial. When A_t_ hits the threshold B, the subject stops investing in favor of starting a new trial. We then constrained A_0_ and B for a model that realized optimal time investment guided by decision confidence. Since, in our task, h(*t*) is an exponential distribution with decay parameter τ, H(*t*) is given by 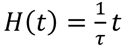 (defined above the lower truncation of the exponential reward delay distribution at 0.5 s disregarding the upper truncation at 8 s, which comprises less than 0.6 % of the distribution’s values) and it follows for optimal time investment T_opt_ when A_t_ hits the threshold B:

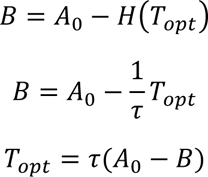

On the other hand, optimal time investment T_opt_ is given by eq. 7 and it follows:

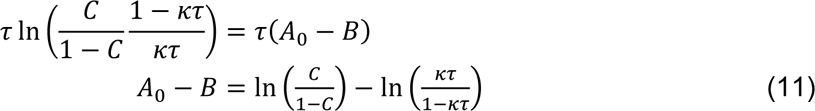

Motivated from the fact that decision confidence is fully determined by the percept of the sensory stimulus and choice, i.e., at the beginning of the waiting period, we set the accumulator’s initial value A_0_ and the threshold B to:

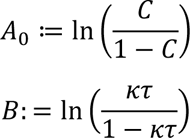

Note that mathematically equivalent notations exist for A_0_ and B if they satisfy eq. 11.

We introduced variability, and therefore sub-optimal time investment, in our model by adding accumulator noise N_acc_(t) (zero-mean Gaussian noise with standard deviation σ_acc_) to A_t_ in each time step t.

To estimate each subject’s magnitude of the accumulator noise we fitted our model using two free parameters, noise magnitude given σ_acc_ and the opportunity cost κ. Treating the opportunity cost of time κ as a free parameter allowed for subject-specific utility functions of time and reward. Note that this approach constrains the shape of the utility function according to eq. 7, while allowing for shifts of this function along the temporal axis. Accumulator noise effectively smoothens this mapping function. For model fits, we minimized the Kullback-Leibler divergence between the empirically observed time investment distribution and the model’s predicted time investment distribution in 1,000,000 trials in a grid search. For κ, grid search comprised values from 0 to 0.15 (0.01 steps); for σ_acc_, we used values from 0 to 0.3 (0.01 steps).

